# Agentic Lab: An Agentic-physical AI system for cell and organoid experimentation and manufacturing

**DOI:** 10.1101/2025.11.11.686354

**Authors:** Wenbo Wang, Simran Swain, Jaeyong Lee, Zuwan Lin, Bradley Canales, Almir Aljović, Yaxuan Liu, Qiang Li, Arnau Marin-Llobet, Mai Liu, Zihan Gao, Ren Liu, Juan R. Alvarez-Dominguez, Jia Liu

## Abstract

Reproducibility in biological research and manufacturing remains constrained by the complexity of multi-step protocols, fragmented data-analysis pipelines, and the intrinsic variability of experimental execution. Here, we present Agentic Lab, an agentic-physical AI platform that unifies large language model and vision language model (LLMs/VLMs)-driven reasoning with real-world laboratory operations. Agentic Lab uses multi-agent orchestration architecture, comprising of specialized subagents for knowledge retrieval, protocol design, multimodal data analysis, and training-free segmentation and representation learning for intrinsically explainable single-cell and organoid phenotyping. These agents operate under the orchestration of a virtual principal investigator MolAgent that is linked to an augmented reality (AR)-based physical AI interface, which can bridge digital reasoning with human physical execution. Agentic Lab perceives real-world experimental activities, provides context-aware instructions, identifying procedural errors in real time for humans to correct, and continuously evolves with its long-term memory database expanding through the accumulation of experimental data logs from human scientists. This interaction allows scientists and AI agents to collaborate and co-evolve dynamically, closing the loop between planning, action, and analysis in the traditional cell and organoid research lifecycle. We demonstrate Agentic Lab in organoid differentiation from human pluripotent stem cells, where it autonomously generates protocols, monitors culture procedures, and identifies subtle morphological heterogeneity linked to growth conditions. The system interprets these phenotypes, grounds them in literature, and proposes targeted instructions for improving differentiation efficiency. By combining multi-agent reasoning with physical laboratory awareness, Agentic Lab transforms experimentation and biomanufacturing from a static workflow into an adaptive, feedback-driven, bidirectional process that integrates agentic AI into the research lifecycle. This framework establishes a foundation for intelligent laboratories that integrate design, execution, and interpretation within a unified agentic-physical system.

## Main

Cell and tissue culture are essential for cell biology research and biomanufacturing. Three-dimensional (3D), stem cell-derived organoid models that replicate human tissue architecture and function *in vitro* have advanced regenerative medicine, cancer therapy, and personalized drug discovery^1,2^. However, organoid generation, maintenance, and downstream assays require complicated coordination of multiple culture steps and experience-based decision-making, necessitating years of training^3^. These challenges are further intensified by the lack of standardized approaches for organoid data collection, analysis, and phenotypic interpretation. Researchers must translate abstract experimental instructions into practical actions, making real-time adjustments guided by visual feedback and personal expertise. The interpretation of data generated through such trial-and-error approaches is subjective (**Fig. 1a**). This results in variable reproducibility, as small deviations in handling or environmental conditions can yield disproportionate effects on organoid development and function, posing a significant layer of uncertainty in cell and organoid research^4^.

**Figure 1.**
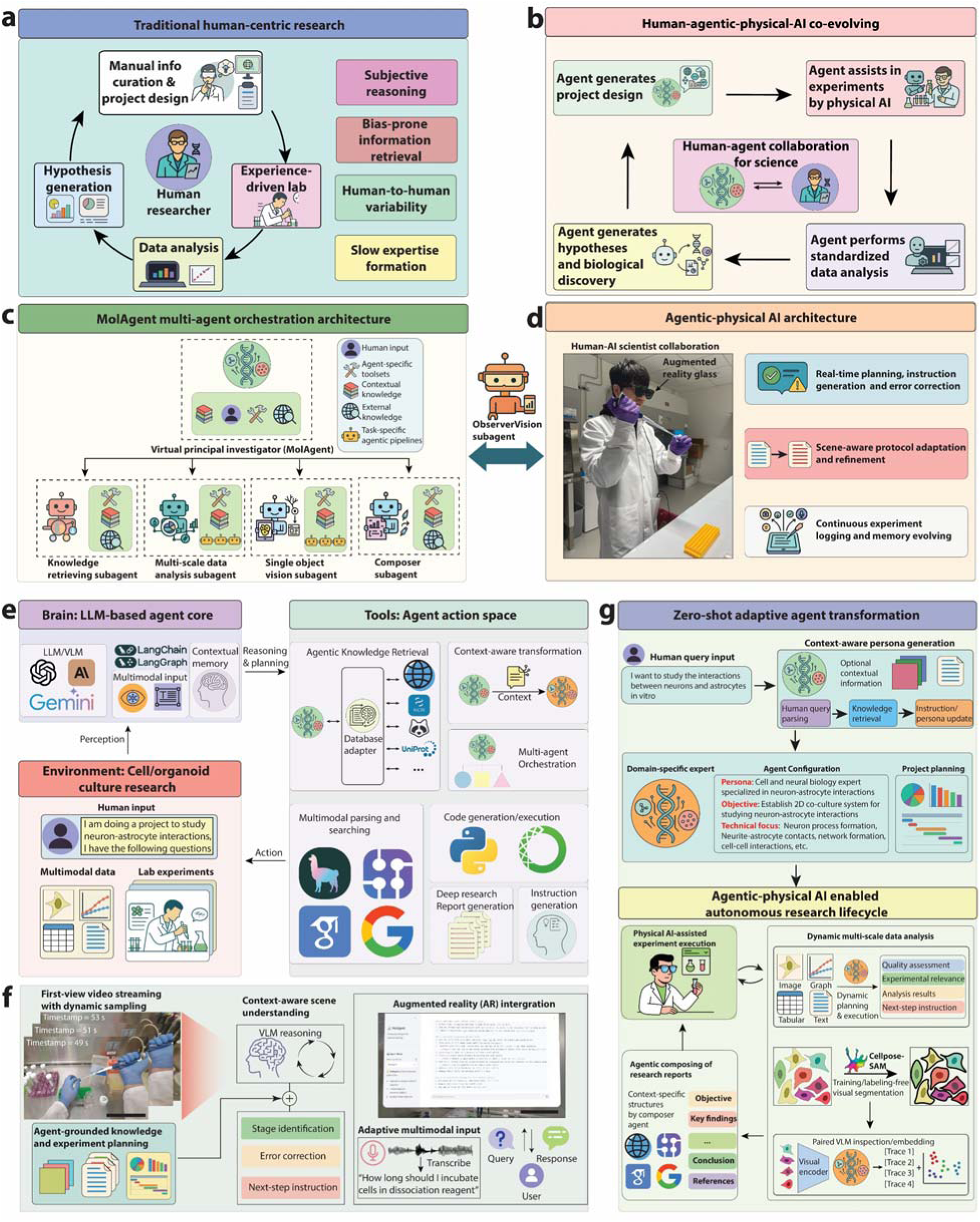
Agentic Lab: An agentic-physical AI platform that unifies multimodal large language models and vision language models (LLMs/VLMs)-driven AI agents with laboratory experimentation via an augmented-reality interface. (a) Traditional human-centric research follows a sequential cycle from hypothesis and experiment to data analysis and reporting, iteratively refining new questions. (b) Human–agentic-physical-AI co-evolving workflow, AR-guided scientists collaborate with autonomous lab agents that generate real-time protocols, analyses, and insights in a continuous closed-loop of discovery. (c-d) Agentic–physical architecture combining a hierarchical multi-agent system with an augmented-reality (AR) interface. A principal-investigator agent coordinates subagents for knowledge retrieval, data analysis, vision, and report composition (c), while the ObserverVision subagent links the AI to the physical lab, enabling real-time guidance, scene-aware protocol adaptation, and continuous experiment logging (d). (e) Core modules integrating multimodal LLM/VLM reasoning, contextual memory, unified tool adapters for retrieval, parsing, and code execution, orchestrated through LangChain/LangGraph. (f) Immersive AR interaction with first-person video streaming, context-aware scene understanding, error correction, and multimodal (voice/text) communication. Scale bar, 200 mm. (g) Zero-shot adaptive agent transformation from user query to domain-specific persona generation and experimental planning, completing a closed-loop workflow from hypothesis to execution and analysis. Photographs in panel (d) are the authors, who have provided consent for their inclusion.

Recent developments in artificial intelligence (AI) and its applications within biological sciences enable robust, high-throughput analysis and interpretations of complex, multimodal data^5–7^. Early AI algorithms have enhanced interrogation of organoid systems by applying statistics and deep learning models for accurate phenotyping that links cell phenotypes with their morphological features and other covariates^8,9^. Natural language-processing (NLP) tools, as another example, aim to improve literature mining and protocol extraction for project planning^10,11^. However, most current tools remain fragmented across computational and experimental domains. For example, vision models can extract low-dimensional representations of individual cells or organoids from images, but these features often lack explainability and contextual relevance. Similarly, literature retrieval tools can identify topic-related references, but they cannot integrate them into actionable, experiment-specific protocols or plans, given the high complexity of experimental planning in cell and organoid biology research. Another challenge is extracting biologically meaningful and human interpretable features from the resulting data. While deep learning models can capture subtle feature differences, mapping these computational features into human understandable labels, such as cell type, differentiation state, or disease status, remains difficult and commonly undergoes manual labeling by experts, the feature and embedding extract cannot be intuitively related to human understandable or biologically plausible feature descriptions^12–14^. Beyond computational limitations, another critical barrier to applying AI to cell biology research and manufacturing is the inherent variability of humans performing experimental procedures. Manual execution can introduce inconsistencies, errors in following instructions, subjective visual assessments, incorrect sample handling, and variable interpretations of protocol instructions within or across laboratories. These execution-level variations lead to difficulties in reproducing results. For example, subtle differences in medium changes, cell passaging timings, or environmental conditions can profoundly impact cell status, and ultimately organoid differentiation or function. Lastly, existing AI systems operate exclusively within the digital realm, unable to perceive or guide the physical laboratory environment while these variations occur, and only in a *post-hoc* manner. Thus, they cannot detect or correct the implementation errors in real-time. Bridging this gap would allow AI systems to extend beyond computational analysis and provide real-time, context-aware support that improves experimental consistency and reproducibility for human scientists.

Recent advances in LLMs and VLMs provide strong foundations for addressing these limitations. These models are pretrained on vast amounts of data across various domains^15,16^. With emergent reasoning capabilities they can interpret protocols, derive knowledge from literature, and analyze multimodal data^17,18^. However, standalone LLMs/VLMs remain largely reactive. They generate outputs when prompted, and lack the capacity to autonomously perceive, plan, or act on the environment^19,20^. Agentic AI addresses this limitation by equipping LLMs/VLMs with an extensible set of tools and action logic that allow it to couple perception, reasoning, and perform intelligent actions on external environments^21,22^. AI agents have been heavily applied to various domains of science including bioinformatics^23–25^, brain machine interfaces^26,27^, behavior analysis^28^, and electronics design^29^. These agents can dynamically interact with experimental data, browse and retrieve external resources, and iteratively reflect their action traces as new evidence emerges. Multi-agent architectures further enhance this capability by delegating tasks among specialized subagents for knowledge retrieval, planning, vision, or analysis^30,31^. The coordinated interactions of subagents mirror the organization in human research teams, supporting parallelization, cross-validation, and integration of new capabilities. Despite this progress, the field still lacks domain-aware AI systems tailored to cell and organoid research. Agentic systems that intelligently retrieve knowledge to generate experimental plans, perform dynamic multimodal data integration, and interpretable phenotypic analysis are needed. Even more critical are systems that can perceive and guide physical laboratory procedures to reduce human execution variability, as current agentic systems remain confined to computational spaces (**Fig. 1b**). They cannot observe laboratory conditions, monitor reagent handling accuracy, detect timing errors, verify confluence assessments in real time, or provide corrective step-by-step instructions during protocol execution. Without direct perception of the lab environment, these systems cannot distinguish whether experimental failures are from biological variance or execution errors, which poses a fundamental limitation for achieving reproducibility. Addressing these gaps requires advanced computational intelligence and physical device interfaces that can extend AI perception into real world laboratory workspaces.

To bridge these digital and physical gaps, we developed Agentic Lab, a context-aware, autonomous agentic-physical AI platform built on a multi-agent architecture specifically designed for cell and organoid research and manufacturing. The central virtual principal investigator (PI) MolAgent delegates tasks to specialized subagents for knowledge retrieval, planning, vision segmentation, multimodal analyses, and synthesis, integrating outputs via a dynamic orchestration layer. Key innovations include: (1) retrieval-augmented generation (RAG) with iterative self-reflection for protocol refinement; (2) augmented-reality (AR) driven physical interaction, where an ObserverVision subagent perceives the laboratory environment through AR glasses to provide context-aware, step-by-step guidance during hands-on execution and real-time monitoring of experimental conditions; (3) dynamic multi-agent planning with feedback-driven revision across imaging, tabular, textual, and graphical modalities; (4) training-free, explainable, morphological phenotyping via paired semantic and quantitative embeddings generated through Cellpose-SAM^32–34^, VLMs and CLIP-based models^35^; and (5) continuous learning that captures human-AI interactions and experimental outcomes to build an evolving knowledge base. We validated Agentic Lab in the differentiation of human pancreatic organoids from human pluripotent stem cells, performing a multi-week protocol in which morphological phenotypes served as critical indicators of differentiation success and batch quality. Agentic Lab autonomously generated an expert-level protocol, assisted human-physical execution through AR-guided monitoring of confluence and timing, identified morphological signatures of batch variability, linked them to culture conditions, and proposed targeted protocol adjustments. By unifying reason-driven computational planning with physical laboratory interfaces, Agentic Lab enhances experimental throughput, transparency, and reproducibility, representing a transformative step towards intelligent, AI-augmented laboratories in organoid biology and beyond.

## Results

### Agentic Lab’s agentic-physical system architecture

The architecture of the Agentic Lab system is designed as a comprehensive agentic-physical collaborator spanning the entire cell and organoid research lifecycle from generating the project design and experiment planning, instruction generation, and data analysis to experimental execution and discovery. The system organization mimics a scientific research group that couples advanced computational reasoning with direct laboratory perception, bridging the gap between digital intelligence and physical experimentation.

At the system’s core is a virtual PI agent MolAgent, which serves as the primary orchestrator, interpreting human instructions, managing experimental context, and coordinating specialized subagents with distinct expertise: a knowledge-retrieving subagent for external literature synthesis through agentic RAG, a multi-scale data analysis agent for processing different types of experimental outputs, a single-object vision agent for deep morphological profiling and generation of intrinsically explainable embeddings, and a composer subagent for creating structured scientific reports **(Fig. 1c)**. To bridge the multi-agentic architecture with physical AI integration, the system incorporates an ObserverVision subagent which incorporates agentic reasoning into the physical laboratory environment. Through first-person video streaming from the augmented-reality (AR) glasses, this subagent perceives experimental procedures in real time with protocol-specific information from the virtual PI MolAgent to deliver context-aware guidance, adapt protocols dynamically, and continuously log experimental progress **(Fig. 1d)**. This modular, hierarchical design allows Agentic Lab to tackle complex research project planning and implementation by decomposing and routing specific tasks to corresponding subagents for processing.

At its LLM/VLM core, MolAgent and its subagents are powered by state-of-the-art multimodal LLMs and VLMs such as GPT-4.1^36^, Claude-Sonnet-4^37^, Gemini 2.5 Pro^38^, etc. The whole agentic system is built upon a dynamic LangChain/LangGraph-based orchestration layer **(Fig. 1e)**^39^. This cognitive engine provides advanced capabilities in reasoning, planning, and perception, equipped with a versatile action space enabling web browsing, multimodal parsing and searching, dynamic code generation and execution, and orchestration of its subagent team by using an agent-as-tool strategy.

The AR-enabled physical layer further extends this agentic intelligence into the physical experiment execution **(Fig. 1f)**. Using AR glasses with first-person video streaming from the front RGB camera, the ObserverVision subagent performs context-aware scene analysis through VLM reasoning to identify protocol stage, detect protocol deviations, and generate corrective instructions directly within the user’s field of view. The interface supports multimodal input like voice and text, allowing seamless dialogue and feedback between the scientist and Agentic Lab during active experimentation **(Fig. 1f)**. This bidirectional human-AI loop ensures that computational guidance remains grounded in the physical reality of the lab-bench, transforming static protocol execution into a dynamic, continuously adaptive collaboration. Moreover, the streamed videos will be stored in the local computation unit, performing dense video parsing and analysis for personalized experimental log recording and summary. Those traces will be stored as long-term memory for the agent to co-evolve with the human scientist to provide more targeted, personalized instructions.

The hybrid agentic-physical system’s primary feature is a zero-shot adaptive logic flow that facilitates dynamic human-agent collaboration and co-evolvement **(Fig. 1g)**. Upon receiving a human query for a project or experimental description such as “*I want to study the interactions between neurons and astrocytes in vitro*”, MolAgent builds and adopts a context-aware persona by retrieving and integrating domain knowledge through routing the request to the knowledge retrieval subagent and collecting knowledge reports from various external databases. The agent transforms itself into a domain-specific expert tailored to the user’s query, for example, becoming a “*Cell and neural biology expert*” specialized in neuron-astrocyte interactions. This allows the system to engage in iterative cycles of experiment planning, real-time physical guidance, dynamic data analysis, and next step instructions that are directly relevant to the specific scientific context.

This complex, adaptive logic is implemented using stateful workflows built upon the LangChain/LangGraph framework (**Extended Data Fig. 1**). The virtual PI MolAgent’s graph encodes the primary decision-making pathways, including whether to engage in standard conversational exchanges, initiate multi-step protocol generations, or other subagents (**Extended Data Fig. 1a**). The specialized subagents, including ObserverVision, operate within their own internal graphs, allowing for modular and robust task execution across both digital and physical domains (**Extended Data Fig. 1b**).

To make these advanced capabilities accessible to scientific researchers, the entire system is available through a simple, web-based frontend (**Extended Data Fig. 2**) based on Streamlit^40^. This interface provides a centralized platform for conversational interaction and agent mode selection, including AR-guided experiment mode (**Extended Data Fig. 2a**), side bar panels for protocol generation and optimization with real-time physical feedback integration (**Extended Data Fig. 2b**), and integrated panels for multimodal data uploading including live video streams from AR glasses, and session history management (**Extended Data Fig. 2c**). This comprehensive and interactive frontend interface makes the powerful agentic-physical backend fully accessible to scientists without requiring computational expertise.

Through this hybrid architecture, Agentic Lab transforms experimental research into a continuously learning, feedback-driven process in which computational intelligence, human expertise, and physical execution evolve together, creating a true closed-loop workflow, from project definition to physical AI-augmented experiment execution, dynamic multi-scale data analysis and discovery generation.

## Multi-agent collaboration for expert-level protocol generation

A core feature of Agentic Lab is its ability to transform high-level user requests into detailed and executable experimental protocols through a sophisticated retrieval-reflection system. This process is initiated by the main virtual PI agent, which leads a multi-agent team to manage the workflow (**Fig. 2a**). Upon receiving a request such as “*I need to develop a protocol to study the interactions between neurons and astrocytes in vitro*”, it first delegates the task of gathering information to its knowledge retrieving subagent. This specialist performs distributed querying across diverse sources, including general web search for broad context, Google Scholar for peer-reviewed literature, and external protocol databases like Protocols.io^41^, UniProt^42^, NCBI^43^, FPbase^44^, etc, through unified API adaptors for established experimental methods. Crucially, rather than processing all retrieved information indiscriminately, the subagent performs an agentic literature grading, critically assessing and ranking the search results for relevance and reliability. It then dynamically parses only the top-ranked sources with a user customizable number of parsing documents, leveraging the agentic parsing engine LlamaParse^45^ to extract and structure unstructured text into digestible knowledge chunks. These parsed chunks are then passed on to the composer subagent. This subagent’s role is to synthesize these inputs into a final, comprehensive knowledge report. The report serves as an external knowledge base, expanding the subagent’s knowledge boundary beyond its internal parametric memory (**Fig. 2b**). We quantitatively benchmarked this subagent’s performance against both vanilla LLMs (prompt only) and LLMs augmented with standard web search capabilities, using a weighted set of criteria designed to assess scientific rigor, comprehensiveness, and practical utility (**Extended Data Fig. 3a**). While this specific agentic approach shows a greater time cost, our results show that the specialized subagent consistently produces significantly higher-quality knowledge bases across multiple LLMs (**Extended Data Fig. 3b-c**). The superior depth and relevance of the retrieved information provide a stronger foundation for subsequent protocol generation. By combining this rich, context-specific external knowledge with its internal knowledge and intrinsic reasoning capabilities, the virtual PI MolAgent is equipped to generate a detailed initial protocol and adopt the specific identity with expertise required for the task (**Fig. 2c**).

**Figure 2.**
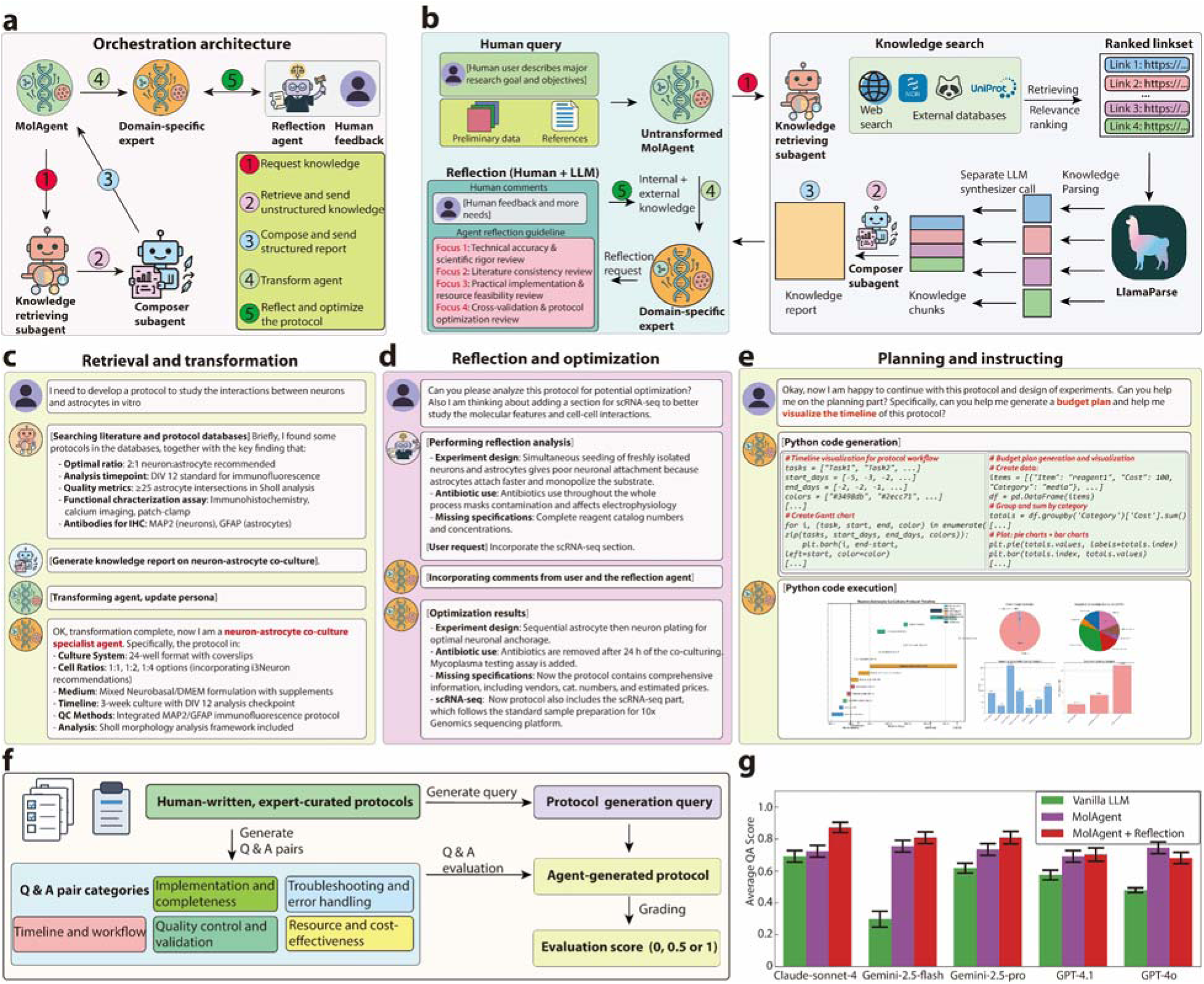
The agentic knowledge retrieval and reflection for adaptive core. **(a)**, The orchestration architecture of the multi-agent system for protocol generation. **(b)**, The human-in-the-loop reflection. **(c)**, Example of the retrieval and transformation, where a user request for a neuron-astrocyte co-culture protocol is processed by the agents to generate a detailed, specific, domain-specific protocol. **(d)**, The reflection and optimization cycle. **(e)**, Examples of dynamic code generation and execution for experiment timeline visualization and budget planning. **(f)**, Evaluation pipeline. Expert-curated, human-written protocols are used to generate question–answer (Q&A) pairs, together with a one-sentence query to prompt vanilla LLM, MolAgent, and MolAgent with reflection for protocol generation. **(g)**, Evaluation results across multiple criteria, showing that MolAgent and MolAgent with reflection outperform vanilla LLM in protocol completeness, quality control, troubleshooting, cost-effectiveness, and workflow accuracy. n = 47 Q&A pairs for each model. Error bars represent mean ± Standard Error of the Mean (SEM).

Following the initial draft, the system implements a rule-based self-reflection logic involving both human input and autonomous reflection with an agent as a judge. This collaborative cycle allows for iterative refinement, where the PI agent can incorporate expert feedback or perform its own analysis to identify and correct potential flaws, such as missing reagent concentrations, ambiguous instructions, or infeasible experimental timelines. For example, in the reflection on the neuron-astrocyte co-culture protocol generation, the judge agent successfully identified as a flaw in the experiment design that the simultaneous seeding of the neurons and astrocytes will lead to poor neuron attachment, as the astrocytes will monopolize the substrate^46^. The agent then provides an updated version to perform sequential seeding of astrocytes and neurons to ensure proper coculture. This process also incorporates the human suggestion to add single cell sequencing (scRNA-seq) experiments to enable better characterization (**Fig. 2d**).

Agentic Lab’s utility extends beyond static text generation by transforming the final protocol into an actionable blueprint for laboratory management through dynamic code generation and execution. Agentic Lab parses the written protocol and translates its instructions into functional planning figures. For instance, it can parse procedural steps to create Gantt chart visualizations that map out the entire experimental schedule, or extract materials lists to generate detailed budgets with itemized cost calculations. These on-the-fly extensions provide researchers with critical tools for logistics and resource management, turning a text-based protocol into comprehensive experimental roadmaps. (**Fig. 2e**, **Extended Data Fig. 4**).

To quantitatively measure the impact of this workflow, we developed a benchmarking framework using expert-derived question-and-answer (Q&A) pairs using LLM to objectively score the completeness and accuracy of generated protocols (**Fig. 2f, Extended Data Fig. 5a**). The results demonstrate that, with more processing time cost (**Extended Data Fig.5b**), Agentic Lab outperforms vanilla LLMs. Furthermore, the inclusion of the autonomous reflection logic consistently yields the highest performance, providing clear evidence that this iterative refinement is critical for generating high-fidelity, expert-level protocols (**Fig. 2g**).

## Multi-modal data analysis through dynamic planning and execution

Modern biological experiments produce a variety of data types from microscopy images to tabular measurements requiring flexible analytical approaches. To address this, Agentic Lab deploys a specialized multi-scale data analysis subagent to handle multimodal data through dynamic planning and execution. This subagent is invoked whenever a user provides experimental data along with an analytical objective. As illustrated in the high-level workflow (**Fig. 3a**), the virtual PI MolAgent first frames the user’s request in the context of the active protocol, ensuring the analysis is informed by the relevant experimental stage. It then dispatches the data and context to the specialized sub-agent. This handoff is critical, as it ensures the multiscale subagent “knows” what experiment the data came from and what questions the researcher is trying to answer.

**Figure 3.**
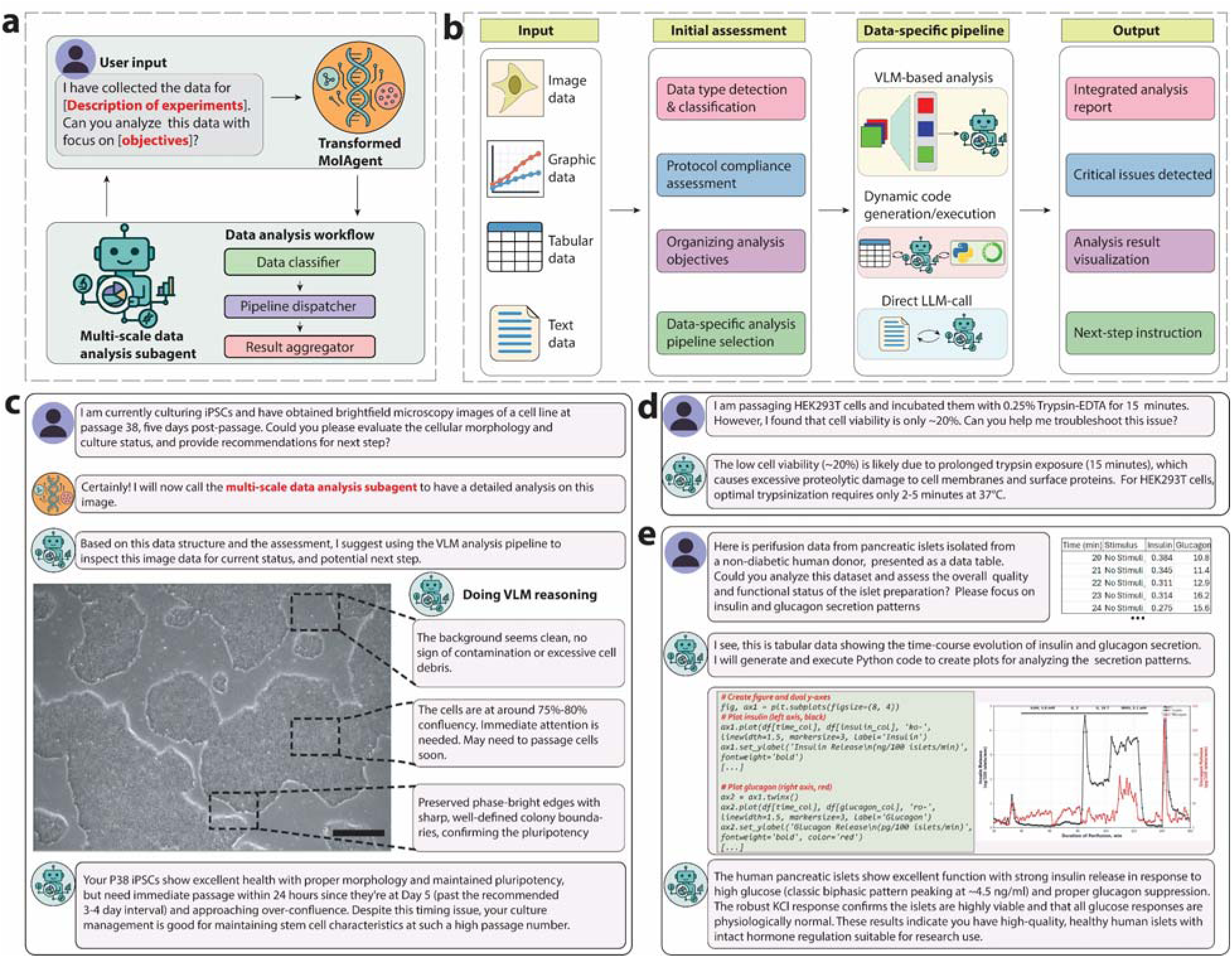
Multimodal data analysis workflow and capabilities of the multi-scale data analysis agent. **(a)**, The high-level data analysis workflow. Upon receiving user input containing experimental data and objectives, the transformed MolAgent invokes the multi-scale data analysis agent, which classifies the data, dispatches it to a specialized pipeline, and aggregates the results for the user. **(b)**, A schematic of the multimodal analysis pipeline. The agent first detects the data type (e.g., image, tabular, text), assesses protocol compliance, and organizes analysis objectives. It then selects a data-specific pipeline, such as VLM-based analysis for images or dynamic code generation for tabular data, to produce an integrated report, visualizations, and next-step instructions. **(c)**, A use case for image data analysis, where the agent applies VLM reasoning to evaluate the morphology, confluency, and health of induced pluripotent stem cells (iPSCs) from a brightfield microscopy image, providing an actionable recommendation. Scale bar, 500 μm. **(d)**, An example of text-based troubleshooting, where the agent diagnoses the cause of low cell viability described by a user and suggests a specific correction to the protocol. **(e)**, A use case for tabular data analysis. The agent processes perfusion data from pancreatic islets, dynamically generates Python code to visualize insulin and glucagon secretion, and interprets the resulting plot to assess the functional quality of the islets.

Internally, the multi-scale data analysis subagent executes a multi-stage reasoning workflow to accommodate diverse data modalities (**Fig. 3b**). It begins with contextualization and planning, where it classifies the data type and, crucially, evaluates it against the active protocol to check for compliance and define analysis goals. Based on this plan, the subagent autonomously selects the most appropriate analysis pipeline. For image data, it leverages a vision-language model (VLM) for sophisticated visual reasoning. For tabular data, it dynamically writes and executes Python code to perform statistical analysis and visualization. Finally, the subagent synthesizes the results into an integrated, human-readable report that includes generated figures, quantitative metrics, and actionable recommendations for the next experimental step.

The versatility of this workflow is demonstrated across several representative cases. For image analysis, the subagent employs its VLM-based pipeline to evaluate iPSC (induced pluripotent stem cell) culture quality, examining morphology and confluency much like an expert biologist. It then integrates its visual assessment with the protocol timeline to provide a specific, context-aware recommendation (**Fig. 3c-e**). For tabular data, the subagent showcases its capacity for autonomous analysis. Given raw data from a pancreatic islet perfusion experiment, it dynamically generates a Python script to create a dual-axis plot of hormone secretion and then interprets the visualization it just created to deliver a high-level scientific conclusion on islet functional quality. The subagent acts as an expert consultant for text-based troubleshooting, diagnosing a user-described problem like low cell viability by cross-referencing it against the protocol and suggesting a concrete solution.

The reliability of this intricate, multi-step workflow hinges on the PI agent’s underlying ability to correctly recognize when and how to use its available tools. To ensure its robustness, we established a rigorous benchmark to quantitatively measure the "tool adherence" of various foundation models serving as the PI agent’s core. We evaluated performance across two critical dimensions: tool usage recognition accuracy (determining *when* a tool is needed) and tool selection accuracy (choosing the *correct* tool for the task) (**Extended Data Fig. 6a-b**). The results reveal that different LLMs exhibit distinct strengths and trade-offs in these agentic capabilities (**Extended Data Fig. 6c**). To determine the optimal overall performer, we developed a weighted performance ranking that prioritizes the crucial decision of when to use a tool. Among the models tested, Claude-sonnet-4 emerged with the highest composite score, driven by its superior accuracy in tool usage recognition at 91.1% (**Extended Data Fig. 6d-e**). This quantitative assessment not only confirms the robustness of Agentic Lab’s decision-making process but also provides an empirical framework for selecting the optimal LLM core to ensure the reliable execution of its multimodal data analysis workflows.

## Single-object analysis enables intrinsic, explainable representations to reveal organoid phenotypes

A central challenge in computational cell biology is the extraction of meaningful, quantitative information from microscopy images. Traditional approaches often rely on either predefined morphometric feature (e.g., size, circularity) which may not capture complex phenotypes, or on deep learning-based embeddings that function as "black boxes." While these numeric embeddings are powerful for classification, their abstract nature obscures the underlying biological meaning, making it difficult for researchers to understand why certain cells are phenotypically distinct. To address this long-standing issue, Agentic Lab introduces an agentic single-object analysis pipeline designed to generate intrinsic, explainable representations of cellular and organoid phenotypes, which is overseen by the SingleObjectVision subagent. We integrate training-free instance segmentation with a reflective optimization loop and a multimodal embedding strategy that aligns visual data with natural language descriptions (**Fig. 4a**). At the core of this strategy is an embedding model, which is uniquely trained to map images and their corresponding text descriptions into a shared, semantically rich, embedding space. By pairing each segmented objects image with a VLM-generated textual description, SingleObjectVision creates an embedding that is not only quantitatively robust but also directly linked to features that are interpretable.

**Figure 4.**
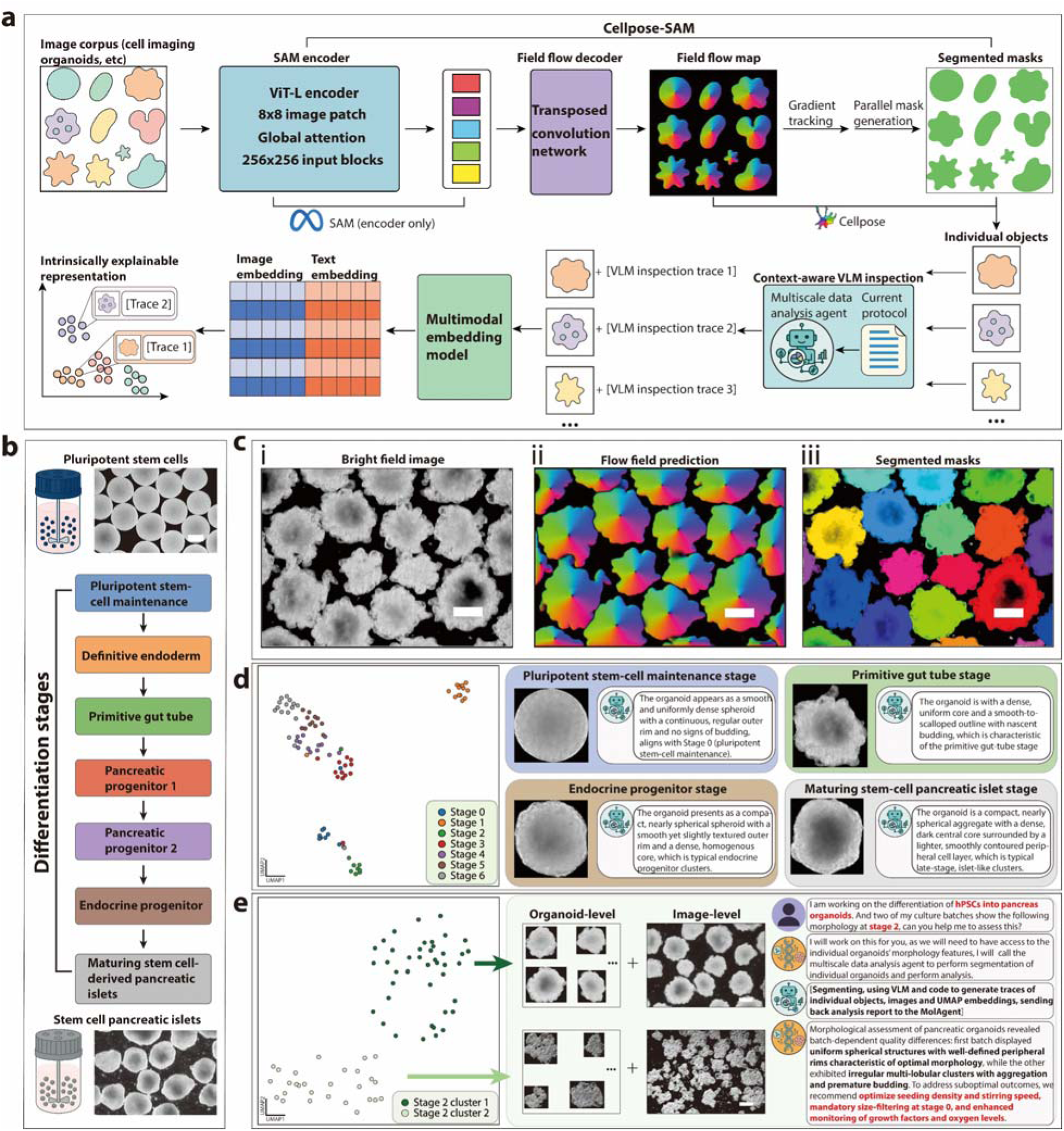
Agentic single-object segmentation and analysis for generating intrinsic, explainable representations. **(a)**, The schematic workflow for integrating Cellpose-SAM, a foundation model for instance segmentation, a VLM to generate descriptive text traces for each object, and a Contrastive Language-Image Pre-training (CLIP)-based models to encode image-text pairs into a joint, explainable multimodal embedding. **(b)**, The differentiation of hPSCs into pancreatic islets is used as the biological model system. **(c)**, Demonstration of the Cellpose-SAM segmentation on a brightfield image of differentiating pancreatic islets at primitive gut tube stage. (i), showing the intermediate flow field prediction (ii) and final instance masks (iii). **(d)**, A UMAP projection of the CLIP embeddings shows distinct clustering of organoids by differentiation stage. Callouts link representative images to their VLM-generated descriptions, highlighting the explainable nature of the representation. **(e)**, A practical use-case where the agent identifies two morphologically distinct sub-clusters within Stage 2. By analyzing the associated VLM descriptions, the agent diagnoses suboptimal culture conditions in one batch and provides actionable recommendations to improve experimental consistency. Scale bars, 200 µm for all panels.

To achieve high-fidelity instance segmentation, we adapted the recently developed Cellpose-SAM^32^, which integrates two state-of-the-art foundation models, Cellpose^34^ and meta’s SAM^33^ model, orchestrating under the dedicated SingleObjectVision subagent. The subagent first leverages Cellpose’s pre-trained model to predict a vector field across the image, where vectors point towards the center of each potential object. From this field, it extracts the coordinates of the vector sinks, which correspond to the predicted object centers. These coordinates are then used as highly informed points that prompt to guide SAM’s powerful, generalizable segmentation engine, generating final and precise instance masks for each individual organoid. This two-stage approach combines the domain-specific expertise of Cellpose in identifying cell-like objects with the zero-shot power of SAM, resulting in a robust segmentation process that requires no experiment-specific training data. To further increase the robustness of the segmentation, we implemented an agentic reflection process that enables iterative self-optimization. The subagent employs vision reasoning with experiment and protocol context to evaluate mask quality, diagnose segmentation errors, and optimize image processing parameters such as the iteration number or resizing of the image to visually aid the segmentation process autonomously, until satisfactory outputs are achieved. The prompt logic underlying this process allows the subagent to inspect, grade, and refine segmentation results through structured reflection. Applied to images of 3D organoids formed during pancreatic differentiation of human pluripotent stem cells (hPSCs)^47^, this self-reflective logic corrected suboptimal Cellpose-SAM outputs and substantially improved the accuracy of cell- and organoid-level masks (**Extended Data Fig. 7)**.

Building on this foundation, SingleObjectVision establishes explainable embeddings through a two-step framework. For every segmented object, a vision–language model generates a descriptive text trace articulating key morphological features in human-readable terms, such as “*a well-formed organoid with smooth, defined borders.*” Each image–description pair is then encoded by a user-selected embedding model, which concatenates them into a shared high-dimensional space that inherently preserves both computational structure and semantic interpretability. As a result, the embeddings provide not only powerful quantitative representations but also intrinsic explanatory value.

We validated the pipeline using the directed differentiation of hPSCs into pancreatic organoids as a model system, where subtle morphological changes are known to be critical indicators of developmental stage and culture quality, but remain difficult to quantify in an interpretable manner (**Fig. 4b**)^47,48^. When embeddings generated by SingleObjectVision were visualized with UMAP, organoids self-organized into distinct clusters corresponding precisely to their known developmental progression (**Fig. 4c-d**). Importantly, the basis for this separation was transparent: representative organoids could be displayed alongside the descriptive text produced by the subagent, allowing each cluster to be understood and rationalized in terms of both visual and semantic features.

Crucially, SingleObjectVision also enabled discovery of unanticipated heterogeneity. As proof-of-concept, organoids from a, nominally, single differentiation stage (Stage 2) were separated into two distinct sub-clusters (**Fig. 4e**). The associated text traces provided direct insight into the morphological basis of this separation: one group consistently exhibited healthy morphology, while the other was described with features such as “irregular shapes with surrounding cellular debris” or “premature clustering.” Based on this interpretable evidence, SingleObjectVision inferred that inconsistent nutrient or growth factor exposure was likely responsible and generated an actionable recommendation to adjust culture conditions. All analysis results are then transferred to the PI MolAgent as an analytical trace, ensuring the full context is available for further interpretation. By grounding its analysis in natural language, Agentic Lab generates hypotheses that could be validated experimentally, effectively bridging sophisticated computational methods with the explanatory requirements of scientific inquiry.

Through its combination of high-fidelity segmentation, reflective self-optimization, and intrinsically interpretable embeddings, Agentic Lab transforms image-based biological analysis into a rigorous, transparent, and actionable framework. This system enables researchers not only to validate prior knowledge of phenotypic progression but also to uncover previously hidden heterogeneity, offering a powerful path toward automated, explainable discovery in cell biology and regenerative medicine.

### Leveraging physical AI for real-time experimental guidance generation

The physical-AI module of Agentic Lab connects the intelligent agentic system directly within the physical lab environment, enabling real-time, context-aware support of cell and organoid culture workflows. Central to this capability are augmented-reality (AR) glasses with 6 degrees of freedom (6 DOF) spatial rendering and anchoring-of-agent user interface, alongside first-person video streaming of laboratory actions through an RGB camera (**Fig. 5a**). The underlying software implementation and context aware scene understanding is embedded in a dedicated subagent, ObserverVision, that directly handles the connection between the physical AI interface and AI agents.

**Figure 5.**
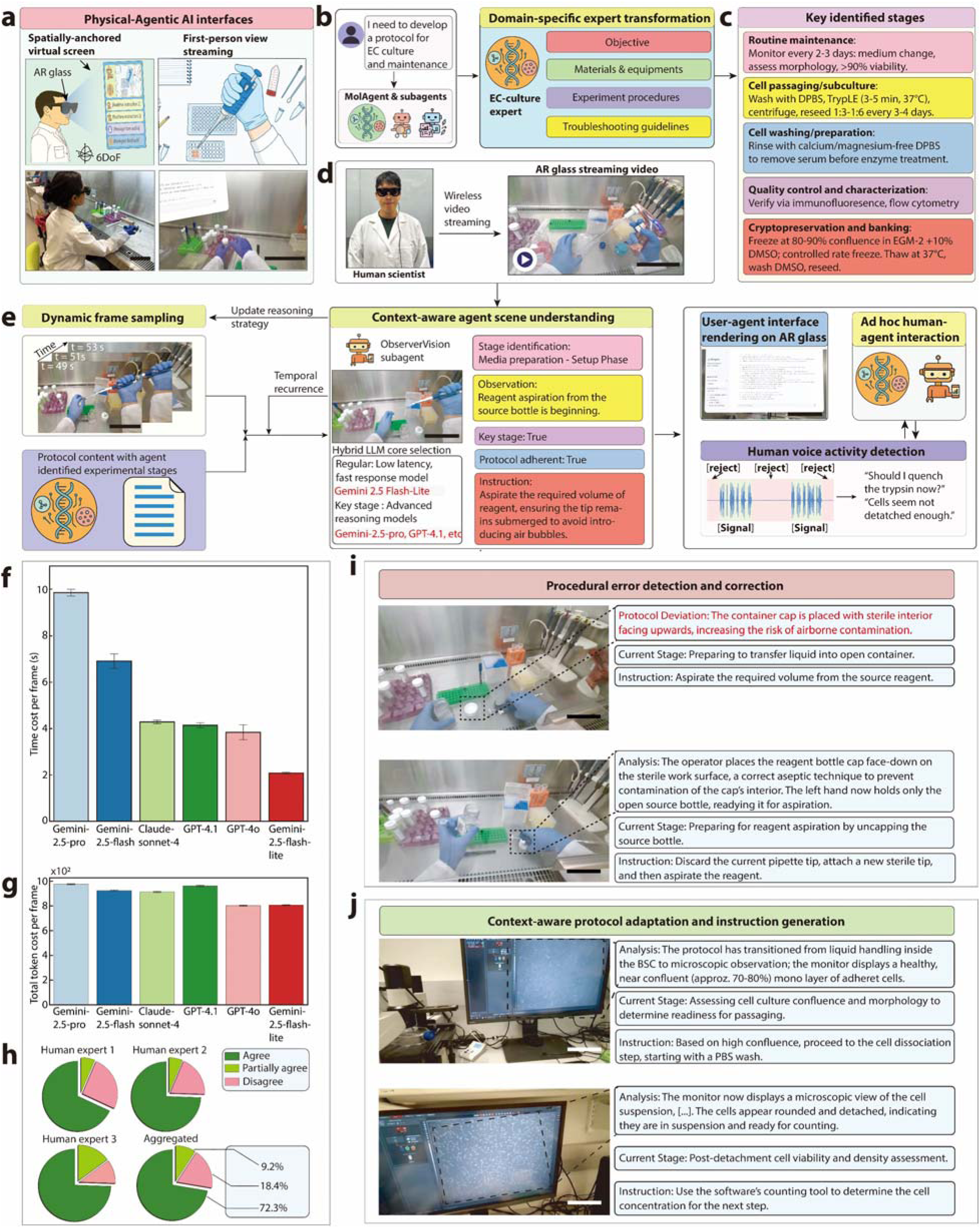
Agentic-physical AI enables seamless human–AI collaboration in cell and organoid biology. **(a)**, The Agentic-physical AI interface integrates AR-based spatial overlays and first-person video streaming, allowing real-time perception of experimental actions. Scale bar, 200 mm. **(b)**, MolAgent transforms into a domain-specific iPSC-culture expert by parsing structured protocol elements including objectives, materials, procedures, and troubleshooting guidelines. **(c)**, The system identifies key experimental stages—media preparation, plate coating, cell revival/thawing, daily maintenance, and passaging—each annotated with corresponding procedural focus points. **(d)**, The AR glasses streaming interface enables closed-loop interaction, where the agent observes the experiment, detects context, and generates adaptive instructions or corrections in real time. Scale bar, 200 mm. **(e)**, ObserverVision subagents perform dynamic frame sampling, multimodal scene understanding, and hybrid LLM-based reasoning for protocol adherence verification and instruction generation. Scale bar, 200 mm. (**f-h**) Benchmarking on time cost per frame (**f**) and total token cost per frame (**g**) of ObserverVision agent for VLM processing capability. n = 633 frames. Error bars represent mean ± SEM. (**h**) Human-expert agreement on instruction generated at key frames identified by ObserverVision subagent. n = 47 cases for instructions. **(i-j)**, Example of procedural error correction (**i**) and instruction generation and protocol adaptation (**j**) during experiment works with physical AI device. Scale bar, 100 mm. Photographs in panel (**a**) and (**d**) are the authors, who have provided consent for their inclusion.

Agentic Lab builds an expert-like experimental persona by interpreting structured protocol data including objectives, materials, procedures, and troubleshooting notes to contextualize its reasoning. Rather than relying on broad laboratory best practices like “*Always wear gloves*”, it grounds its understanding in the specific procedural logic of each experimental stage. (**Fig. 5b**). By coupling the reasoning models with real-time video input, the subagent can recognize which stage of the experiment is in progress such as media preparation, plate-coating, cell thawing, routine cell culture maintenance or passaging and anticipate the procedural focus for each stage (**Fig. 5c**).

Once the workflow stage is identified and the operational context is established, ObserverVision initiates a closed-loop collaboration with the scientist. Through an augmented reality (AR) interface, the system delivers adaptive, real-time instructions and corrective prompts that guide experimental execution. Bidirectional communication between the human operator, MolAgent, and ObserverVision is enabled through multimodal text and dynamic audio channels, allowing direct communication with the subagent through audio input (**Fig. 5d-e**). This design ensures seamless human–physical AI interaction: the scientist retains full control over the experiment setup, while the subagent observes the environment, reasons for ongoing procedures, and provides context-aware guidance precisely at the point of action.

A central engineering challenge in implementing this capability is latency in visual–language reasoning and instruction generation. Larger VLMs offer deeper reasoning and richer scene understanding but incur longer response times that disrupt real-time communication. To balance these trade-offs, we developed a hybrid agent core and dynamic frame-sampling mechanism. During routine monitoring, a lightweight, fast-response VLM core ensures real-time feedback. When a key frame indicating a critical experimental step is detected, the system dynamically invokes an advanced reasoning engine that engages larger VLMs for comprehensive scene interpretation and instruction synthesis, adapting latency to task complexity.

We benchmarked model performance across multiple models and selected Gemini 2.5 Flash Lite as the fast-track model owing to its lowest latency (2.08 ± 0.02 s per 1080p frame). For critical reasoning tasks, we employed advanced reasoning models like Gemini 2.5 Pro, GPT-4.1, Claude-Sonnet-4, etc., for their superior multimodal understanding and instruction quality (**Fig. 5f-g**). To assess instruction reliability, we compared ObserverVision-generated guidance with expert annotations on in-house generated cell culture videos recorded by experimentalists. Across all evaluated key frames, 72.3% of ObserverVision-generated instructions were in full agreement with expert consensus, and an additional 9.2% were partially aligned, indicating over 80% overall concordance between the Agentic Lab system and human scientists (**Fig. 5h**).

As proof of concept, we used the physical AI device to guide and instruct the passaging process of human induced pluripotent stem cell-derived endothelial cells (iPSC-ECs). ObserverVision successfully identified procedural errors, such as placing the container cap facing upward, which increases the risk of airborne contamination, and flagging them as protocol deviations, prompting the human scientist to correct them (**Fig. 5i**). Another example highlights the subagent’s ability to adapt protocols: rather than rigidly adhering to the specified dissociation reagent incubation time, the subagent reasoned based on microscopic images displayed in the user interface to determine whether the cells were sufficiently confluent for passaging and whether the dissociation process had reached the appropriate incubation stage (**Fig. 5j**).

Together, these components enable Agentic Lab’s physical-AI module to convert the standard laboratory environment into a cognitively augmented environment in which experiment execution, monitoring and adjustments are integrated. The researcher maintains oversight and decision-making, but within a tightly coupled human–agent loop that elevates reproducibility, protocol adherence, and experimental agility.

## Multi-agent collaboration enables human-agentic-physical-AI continuous co-evolvement and paper-like research report generation

A defining capability of Agentic Lab lies in its capacity to continuously learn from the complete history of human–agent collaboration and to transform these experiences into structured, scientifically relevant knowledge. Each experimental session produces a comprehensive set of interaction traces including curated external knowledge, reasoning pathways, multimodal data analyses, and intrinsically explainable visual representations that together constitute the agent’s evolving learning record (**Fig. 6a, left**). These traces, enriched by contextual information from the physical-AI interface such as AR-derived video, agent generated instructions, and agent-human interaction logs, are parsed post-experiment to extract temporally aligned behavioral and procedural information (**Fig. 6a, right**). This closed-loop design enables Agentic Lab to not merely document experimental activity but to interpret and internalize it as part of a growing long-term memory database.

**Figure 6.**
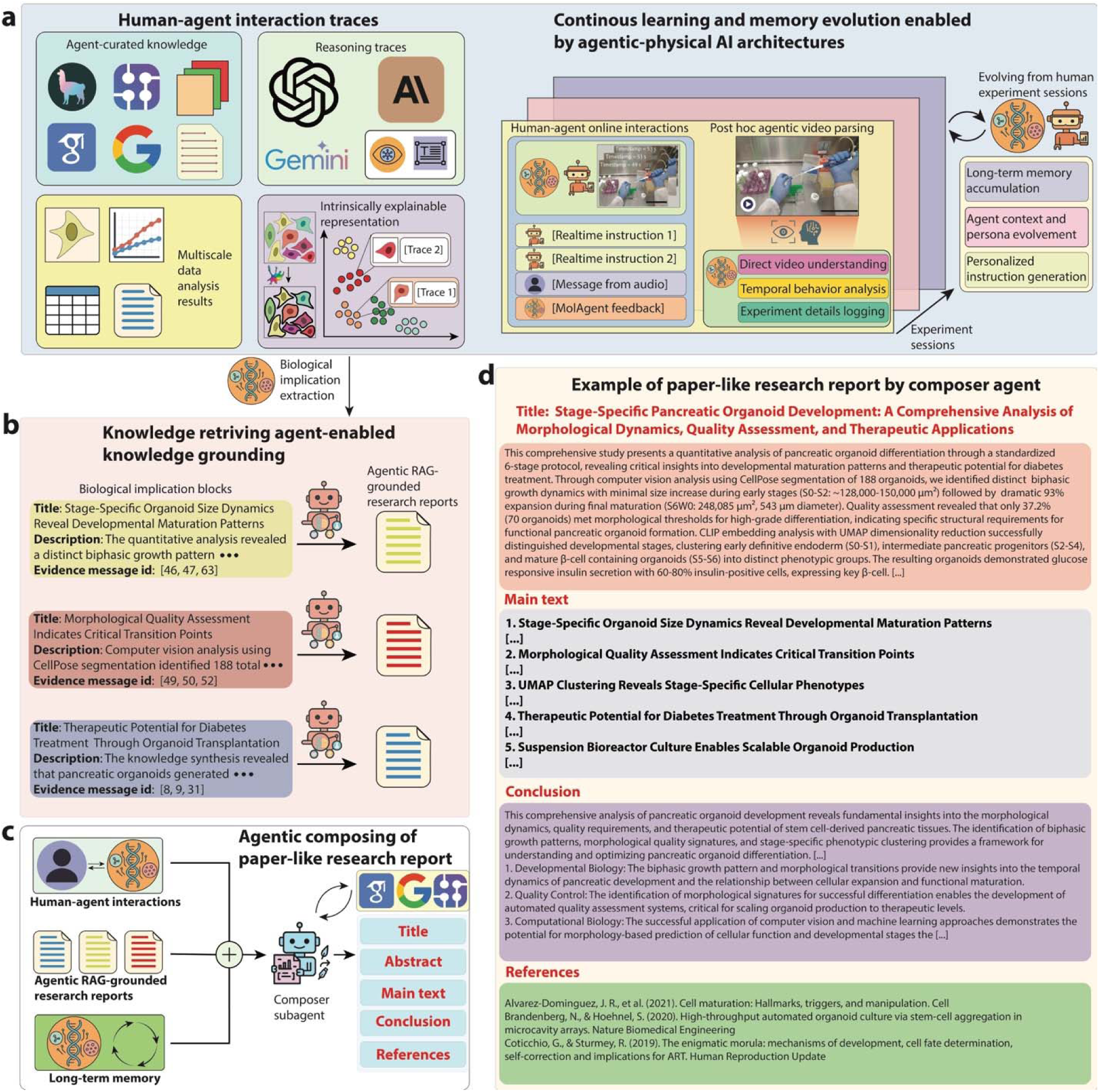
Continuous learning and memory evolution in Agentic Lab system. **(a)** Human–agent interaction traces, curated knowledge, multimodal analyses, and explainable visualizations, together with physical AI interactions, evolve into digital experiment logs that refine future actions and improve human-agent collaboration and experimentation. Scale bar, 200 mm. **(b)** From these traces, the system extracts biological implication blocks that summarize experiment-derived findings. Each block is validated and expanded through targeted knowledge retrieval, generating grounded scientific evidence for integration into Agentic Lab’s internal memory. **(c)** Schematic for the report generation. The Composer agent synthesizes these grounded implications with human–agent logs into structured, paper-like research reports containing title, abstract, results, conclusions, and references. These documents serve both as human-readable outputs and as machine-readable knowledge objects for future reasoning. **(d)** Example of a report generated from Agentic Lab.

From these accumulated traces, the system identifies and structures biological implication blocks that encapsulate experiment-derived insights (**Fig. 6b**). Each block, representing a finding such as stage-specific morphological dynamics or critical transition points in organoid development, is validated and expanded through targeted RAG. By querying relevant literature, the knowledge-retrieving subagent enriches these internal discoveries with external scientific evidence, linking each block to its supporting data via traceable identifiers. This process grounds Agentic Lab’s experiential knowledge in peer-reviewed context, ensuring interpretability and reproducibility.

The composer subagent then synthesizes these grounded implications with the experimental logs into comprehensive, paper-like research reports (**Fig. 6c-d**). Each report includes a title, abstract, main text, conclusion, and reference list, organized according to the thematic structure of the derived implication blocks. These reports serve a dual function: they are human-readable outputs suitable for scientific communication and simultaneously machine-readable knowledge objects integrated into Agentic Lab’s long-term memory. Over successive sessions, these documents accumulate into a dynamic and evolving internal knowledge database.

This architecture supports a self-improving learning loop, in which each experimental interaction refines the PI agent’s procedural memory, contextual understanding, and persona specialization. As the Agentic Lab analyses real-time sensory inputs from the physical-AI layer such as AR video streams, human voice commands, and execution logs, it updates its long-term memory databases and personalized instruction strategies. The resulting memory evolution allows ObserverVision to adapt its future guidance and recommendations to user preferences and laboratory conditions, progressively enhancing experimental accuracy and efficiency. This continuous learning framework transforms Agentic Lab from a static reasoning engine into a self-evolving collaborator that unites digital cognition with embodied scientific experience.

## Discussion

Agentic Lab marks a major progress toward realizing agentic AI systems that closely mimic the organization of human research teams. Its multi-agent architecture recapitulates the way real scientists formulate research projects, plan experiments, interpret data, refine protocols, and generate biological hypotheses. Crucially, Agentic Lab generates intrinsic explanatory embeddings for single-object phenotypes, enabling interpretable mappings from imaging observations to infer biological states. These embeddings not only support downstream classification or clustering but allow users to trace which visual features (e.g. morphology, texture, spatial distribution) drove interpretations. By combining this capacity with knowledge retrieval, reflective planning loops, multimodal analysis, and physical AI instructing under a coherent framework, Agentic Lab spans the full experimental lifecycle from protocol design through data interpretation to hypothesis generation in a unified continuum. This multi-agent architecture addresses a key limitation of many pipelines, which tend to decouple data model training from experiment planning and thus lack interpretability. In contrast, Agentic Lab’s reasoning-oriented core allows it to generalize for unseen protocols, diverse data types, and novel experimental questions, moving us towards adaptive scientific assistants.

Nevertheless, Agentic Lab currently faces several limitations, which we acknowledge as opportunities for future advancement. First, our demonstrations so far are confined mostly to brightfield imaging of organoid phenotypes; applying this system to fluorescence microscopy, multiplexed immunostaining, volumetric imaging, and time-lapse live-cell imaging would further demonstrate its data analysis capabilities. By incorporating visual grounding, spatial-temporal segmentation, and 3D promotable modules, Agentic Lab can learn to parse volumetric or temporal phenotypes for further phenotypic interpretation. Second, Agentic Lab’s memory and domain knowledge currently relies on retrieval from one-time external databases and internal context windows, which limits its ability to retain experimental history or biological insight over long experiments. Embedding a graph-based memory layer or fine-tuning open models via low-rank adaptation (LoRA)^49^ to update parametric memory would allow Agentic Lab to internalize and recall relationships, past experiment outcomes, or lab-specific domain rules. The recent Memory-R1^50^ framework, which uses reinforcement learning to train agents that manage external memory and filter relevant context, offers a promising direction for memory-aware agents. Third, while object-level segmentation would enhance agent decision-making by providing explicit information about reagents and equipment, current approaches face two key challenges: (1) computational overhead of dense segmentation adds latency that may exceed latency budget; and (2) general-purpose models lack robustness to the diverse, specialized equipment in biological laboratories. While lightweight network architectures such as YOLO^51^ or FastSAM^52^ show promise in specific domains, achieving reliable real-time segmentation across biological workflows remains an open challenge. Finally, the self-reflection loops in the current Agentic Lab are largely rule-based or rely on VLM checks, which constrain adaptive learning. A potential direction is to formalize protocol generation, reflection, experiment planning, and data analysis as sequential decision-making problems. These problems are amenable to reinforcement learning (RL) techniques such as Proximal Policy Optimization (PPO)^53^ or Group Relative Policy Optimization (GRPO)^54^, which have already been used to fine-tune LLMs. These methods could allow Agentic Lab to adaptively refine its strategies, learn from successes and failures, and evolve autonomously over time.

Looking ahead, Agentic Lab establishes a foundation for the next generation of intelligent laboratory environments in which agentic reasoning, physical execution and lifelong learning converge. With its implemented AR interface, Agentic Lab already demonstrates how an agentic system can perceive bench procedures in real time, provide context-aware guidance, flag deviations and automatically document experimental metadata, marking a transition from computational assistance to embodied collaboration, where human scientists and agents share the same physical workspace. Building on this foundation, the next frontier lies in coupling Agentic Lab’s reasoning core with vision-language-action (VLA) models^55^, robot foundation models^56^, and general-purpose robotic platforms capable of executing the full experimental workflow, from sample preparation and liquid handling to organoid culture, imaging and analysis. Advances in embodied AI, modular manipulators and immersive perception are making this vision increasingly feasible, as general-purpose robots emerge that adapt dynamically to diverse experimental tasks rather than being confined to fixed action spaces in current robotics for science paradigms^57–59^. Within this framework, interaction traces from human-AI-robot experiments continuously refine Agentic Lab’s semantic, procedural, and episodic memories, enabling its memory layers to co-evolve with robotic actuation. Over time, this transformation turns Agentic Lab from an interactive collaborator into a fully autonomous discovery engine capable of designing, executing, and analyzing experiments from end-to-end.

While our demonstration focuses on organoid and cellular biology, the design of Agentic Lab is intentionally domain-agnostic. The paradigm of agentic orchestration, explainable reasoning, iterative self-evolving, and physical AI integration is applicable to materials science, synthetic biology, electronics fabrication, neuroengineering, and other experimental domains. As these capabilities evolve, platforms like Agentic Lab may reshape the role of scientific researchers, shifting effort away from manual execution and data wrangling, towards conceptual formulation, experimental design, and ethical supervision. By reducing the technical burden of experimentation while preserving human judgment, these agents and physical AI systems have the potential to accelerate scientific innovation and democratize access to state-of-the-art discoveries across institutions worldwide.

## Acknowledgements

J.Liu acknowledges the support from NIH/NIDDK 1DP1DK130673 and NIH/NLM 5R01LM014465. J.R.A-D. acknowledges the support from the Human Islet Research Network (U24DK104162), NIH/NIGMS R35GM157320, Breakthrough T1D (INO20 2025-1707-A-N) and a pilot award from the Diabetes Research Center at the University of Pennsylvania (P30DK19525). J.R.A-D., was also supported by grants to Douglas A. Melton from the JDRF (COE-2020-967-M-N), and the JPB Foundation (award no. 1094).

## Author contributions

W.W. and J.Liu conceived the study. W.W. designed and implemented the Agentic Lab framework, developed both backend and frontend methodologies. S.S. developed the interface for single-object segmentation and physical AI and contributed to tool implementation. W.W., S.S., and Y.L. performed benchmark tests. Z.L. and J. Lee provided critical feedback and discussion throughout the development of Agentic Lab. W.W., B.C., M.L., Z.G., and R.L. cultured the cells used in the study. A.A., Q.L., A.ML, and J.R.A-D. discussed the results. W.W., S.S., and B.C. prepared the figures. W.W. drafted the manuscript. All authors contributed to manuscript discussions and refinement. J.Liu provided overall supervision.

## Methods

### Agentic Lab framework

The backend infrastructure of Agentic Lab was developed using LangGraph, which provides a state-machine framework for building agentic workflows driven by large language models (LLMs). LangGraph enables structured control flow, state persistence, and conditional routing across multi-step reasoning tasks, allowing the system to manage complex experimental design and protocol generation workflows. The system integrates multimodal LLMs from OpenAI, Anthropic, and google Gemini, selected for their advanced reasoning, tool-calling, and vision–language capabilities. These models were used without fine-tuning to preserve their general-purpose adaptability across biological and chemical experimental domains.

The architecture implements a supervisor scheme where the virtual PI agent, MolAgent, functions as the central orchestrator and coordinator, while specialized subagents handle domain-specific tasks. Each agent and subagent preserves its own control logic, system prompt, and toolset, enabling modular and specialized functionality. MolAgent parses user queries, decomposes them into executable tasks, and delegates them to specific subagents for execution based on task requirements. Communication between MolAgent and subagents follows an agent-as-tool strategy that ensures well-defined agent-to-agent interfaces. Agent workflows follow a reasoning-action (ReAct)^60^ loop: (1) the agent receives a user query or experimental request with contextual inputs, (2) selects and invokes appropriate subagents or tools for knowledge retrieval, analysis, internet searching, or protocol generation, (3) synthesizes information and generates structured outputs, and (4) returns protocols, visualizations, or analysis results for user review or further refinement. Tasks such as literature search, data analysis, protocol synthesis, and experimental timeline generation are handled through dynamically selected tools or generated code, executed in real time using an isolated Python environment. The platform supports memory-aware interactions through structured state management, enabling continuity across complex multi-step workflows while implementing safeguards to prevent infinite loops and context overflow.

## Agentic knowledge retrieval

Agentic knowledge retrieval is implemented through a specialized subagent based on the agentic RAG paradigm to enable deep-research capabilities. Upon receiving a formatted query from MolAgent, the subagent generates 5-8 strategic search subqueries and selects appropriate tools from available resources including protocols.io, PubMed^43^, FPbase^44^, Google Scholar, and general web search. The subagent then performs parallel tool execution with quality scoring that weights source reliability, publication date, content length, and keyword relevance to prioritize the most valuable information sources. Prioritized references are parsed through LlamaParse for agentic summarization into knowledge chunks that extract methodologies, numerical data, biological entities, and key findings while preserving scientific terminology. The knowledge chunks and additional search results are then synthesized into a comprehensive knowledge report with source citations, which is returned to MolAgent for downstream task execution or user review.

## Protocol generation, reflection and agent adaptive core

At the center of Agentic Lab is its adaptive core that automatically specializes experimental protocols based on user requests by integrating the internal parametric memory of LLMs with external knowledge retrieved through specialized subagents. Upon receiving a query, for example, "*Generate a protocol for neuron-astrocyte co-culture*", MolAgent formulates a structured research request that specifies experimental objectives, required methodologies, and domain-specific constraints. This formulated query is dispatched to the Knowledge Retrieving subagent, which performs comprehensive multi-source research and returns knowledge. The generated knowledge reports are then transmitted to the Composer subagent, which synthesizes the retrieved information into a structured experimental protocol. The protocol is subsequently optimized through a rule-based reflection process, where an agent-as-judge evaluates the protocol based on technical accuracy and scientific rigor, consistency with current literature, experimental feasibility, and cross-validation with online sources, with optional human queries to address specific experimental demands or constraints. The optimized protocol transforms the canonical MolAgent into a domain-specific expert agent for that experimental system, and the designation is stored in the conversation context and drives subsequent interactions, including project timeline generation, reagent budget planning, experimental design optimization, and real-time guidance during protocol execution through the ObserverVision subagent.

## Dynamic code generation, execution and data analysis

Agentic Lab integrates a Python code execution environment to enable dynamic code and execution. The Python REPL (Read-Eval-Print Loop) executor operates as an isolated tool within the LangGraph framework, allowing the LLM to generate and execute arbitrary Python code. The execution environment maintains isolated local scope with controlled global context to prevent cross-contamination between executions, while automatically capturing standard output, error messages, and Matplotlib-generated figures without requiring explicit saving commands from the LLM. Generated visualizations are automatically encoded in base64 format for inline display, with comprehensive exception handling providing formatted tracebacks to enable iterative debugging. This architecture supports tasks ranging from simple to complex machine learning inference and multi-panel figure generation.

For comprehensive interpretation of heterogeneous experimental data, Agentic Lab implements a Multi-scale Data Analysis subagent for intelligent analysis across multiple data modalities. The subagent automatically detects uploaded files from session state and performs data type classification (image, tabular, or text), then routes each data item to appropriate specialized analyzers: a vision-language model interpreter for microscopy images and experimental photographs, a text interpreter for extracting structured information from protocols and laboratory notes, and a tabular analyzer for statistical analysis of CSV and Excel datasets. The analysis workflow consists of three sequential stages: analyzer selection based on data type and query requirements, parallel execution of individual analyzers with protocol-aware interpretation, and cross-modal synthesis where an LLM aggregates findings across all data sources to identify overarching patterns, contradictions, and research implications. Each analyzer generates structured results including interpretation text, confidence scores, key findings, and optional visualization overlays, which are saved to persistent storage with timestamp and metadata for reproducibility.

## Intrinsic explainable embeddings generation

The SingleObjectVision subagent implements an automated agentic pipeline for generating interpretable, language-grounded representations of single-cell or organoid morphology through integration of instance segmentation, vision-language description, and multimodal embedding. Individual objects are extracted from microscopy images using CellPose with flow field-based boundary detection, followed by automated VLM-based quality reflection for segmentation errors; poor-quality mask segmentation triggers image preprocessing and re-execution with adjusted parameters determined by SingleObjectVision subagent. Segmented objects undergo morphometric filtering based on circularity and physical area derived from scale bar detection, with VLM assessment of biological relevance using parallel asynchronous processing. For each retained object, the VLM generates protocol-aware natural language descriptions capturing morphological features (size, shape, texture, boundary characteristics), which are then encoded alongside object images using the available multimodal embedding models; CLIP^35^ (ViT-g-14, 1,024-dimensional image and text embeddings), SigLip^61^, SigLip 2^62^, or DinoV2 with dino.txt^63,64^. Text and image embeddings are concatenated into joint representations, subjected to principal component analysis and UMAP dimensionality reduction using Scanpy, and formatted as AnnData objects storing embeddings, metadata, and UMAP coordinates for downstream visualization.

## Physical AI configuration and agentic-physical AI integration

The ObserverVision subagent enables real-time augmented reality guidance through integration with XREAL ONE PRO AR glasses equipped with an RGB camera (60 frames per second) with six-degrees-of-freedom (6DOF) inertial measurement unit sensors. The system architecture consists of innate operation system for camera capture and AR rendering, an asynchronous Python WebSocket server for bidirectional communication, and the ObserverVision subagent implemented as a LangGraph workflow for continuous frame analysis, failure mode detection, and instruction generation. Camera frames are transmitted via WebSocket with user defined subsampling frame, and original frame at 60 FPS. Each frame undergoes protocol-aware analysis through ObserverVision subagent with dynamically injected system prompts incorporating the current experimental protocol context retrieved from session state.

Each received frame was processed by the ObserverVision subagent, which dynamically constructed by retrieving the current experimental protocol from Streamlit session state and then identified key experimental stages, for example, “*media preparation, plate coating, cell revival and thawing, daily maintenance, passaging*”, with corresponding procedural focus points. ObserverVision generated concise 2-3 sentence guidance responses for regular frames, which were formatted as AR text overlays with position anchoring. We explicitly use the Gemini series models like Gemini 2.5 pro and Gemini flash lite for its superior performance in image and video understanding capabilities.

Critical moments requiring detailed video analysis were identified using a hybrid keyframe detection algorithm that combined LLM-based keyword extraction from ObserverVision responses with protocol-specific trigger rules. When a keyframe was detected, the WebSocket server transmitted a recording command to the local computing client, which initiated capture of a 5-second video segment at 60 frames per second. The recorded video was uploaded sequentially to the backend server and forwarded to the main MolAgent agent graph for comprehensive analysis including technique validation, protocol compliance assessment, safety verification, and generation of detailed procedural guidance.

## Memory evolution from experimental traces

The agentic-physical architecture enables continuous learning through real-time parsing of augmented reality video streams, audio commands, and laboratory observations during successive experimental sessions, updating temporal behavior analysis, long-term memory stores, and context-aware agent personas. Memory is maintained across three hierarchical levels: ephemeral session state tracking active experimental context, persistent session records serialized as timestamped JSON files with complete experimental provenance, and a cross-session knowledge base caching parsed documents, embedding vectors, and protocol libraries indexed for rapid retrieval. Accumulated traces evolve into personalized digital experiment logs that inform future protocol recommendations and experimental guidance, establishing a self-updating feedback loop between human collaboration, physical experimentation, and autonomous scientific reasoning. This architecture enables temporal tracking of experimental evolution, cross-experiment knowledge transfer through session reloading, and laboratory-wide knowledge aggregation via semantic search across all recorded sessions.

## Frontend of the Agentic Lab

The Agentic Lab user interface was implemented using Streamlit, a Python-based web framework providing browser-based access to multi-agent workflows through centralized session state management. The interface enables dynamic switching between three agent modes: MolAgent for general biological research assistance, SingleObjectVision for morphological analysis with configurable embedding models and dimensionality reduction parameters, and ObserverVision for real-time augmented reality guidance via WebSocket integration with XREAL glasses. Users upload context files including microscopy images and protocol documents through a multi-file uploader with automatic document parsing via LlamaParse, while protocol selection and agent specialization inject domain-specific context into system prompts to generate expert personas. The chat interface renders conversation histories with LangChain message objects, displaying user queries, agent responses with expandable tool call details, and multimodal content including text and base64-encoded images with automatic size management. Visualizations generated during analysis are automatically captured and displayed with associated metadata, while Python code execution outputs are rendered with syntax highlighting. Complete chat sessions including conversation context, agent configurations, analysis results, and embedded visualizations are serialized to JSON format for persistent storage and cross-session reproducibility, enabling temporal tracking of experimental evolution and laboratory-wide knowledge aggregation through session loading and semantic search capabilities.

## Code and data availability

Code and data will be made publicly available at https://github.com/LiuLab-Bioelectronics-Harvard/Agentic_lab upon publication.

---

**Extended Figure 1.**
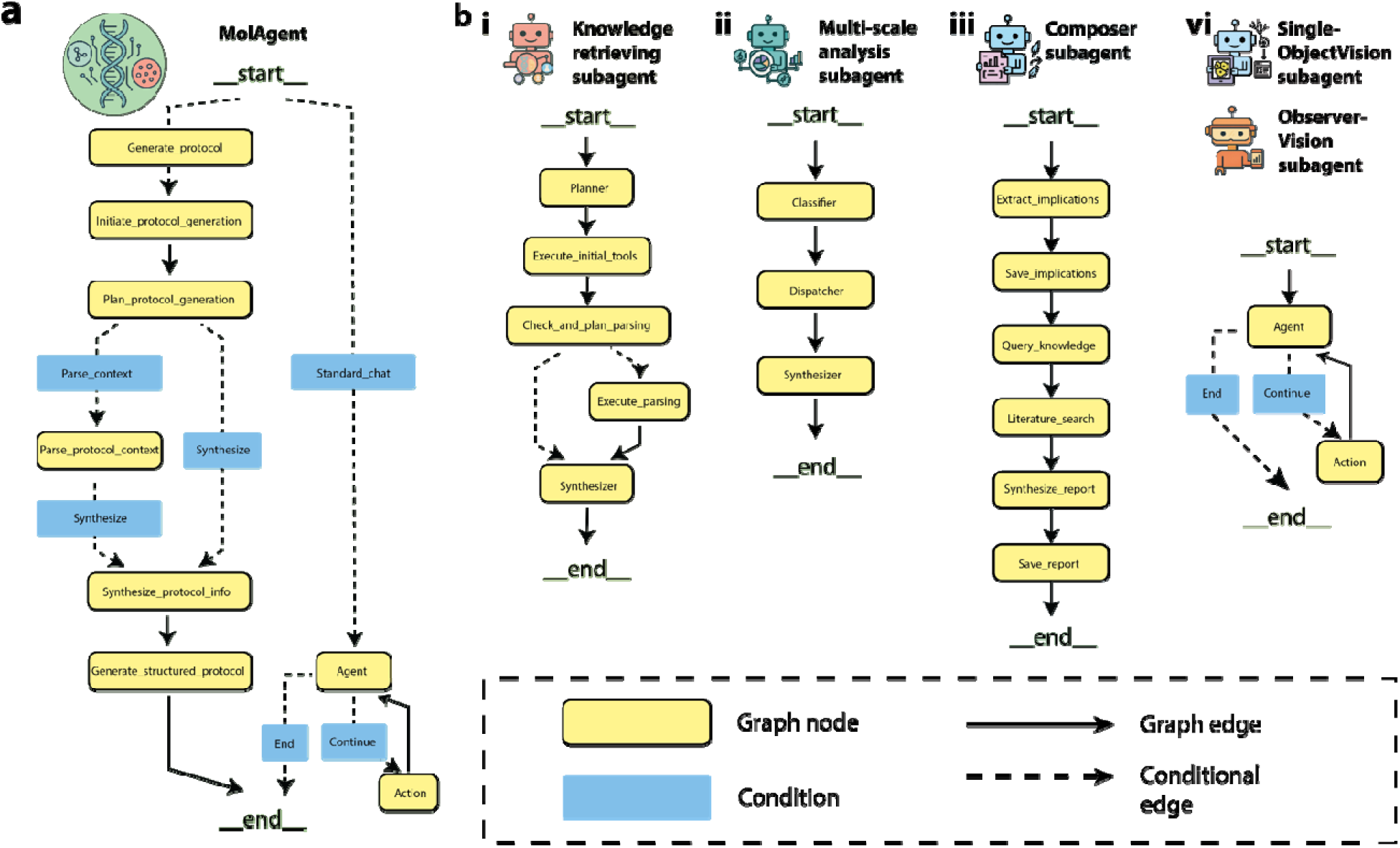
LangChain/LangGraph agent graph architectures in Agentic Lab. (a-b), Agent graph of MolAgent (a), and specialized subagents (b).

**Extended Figure 2.**
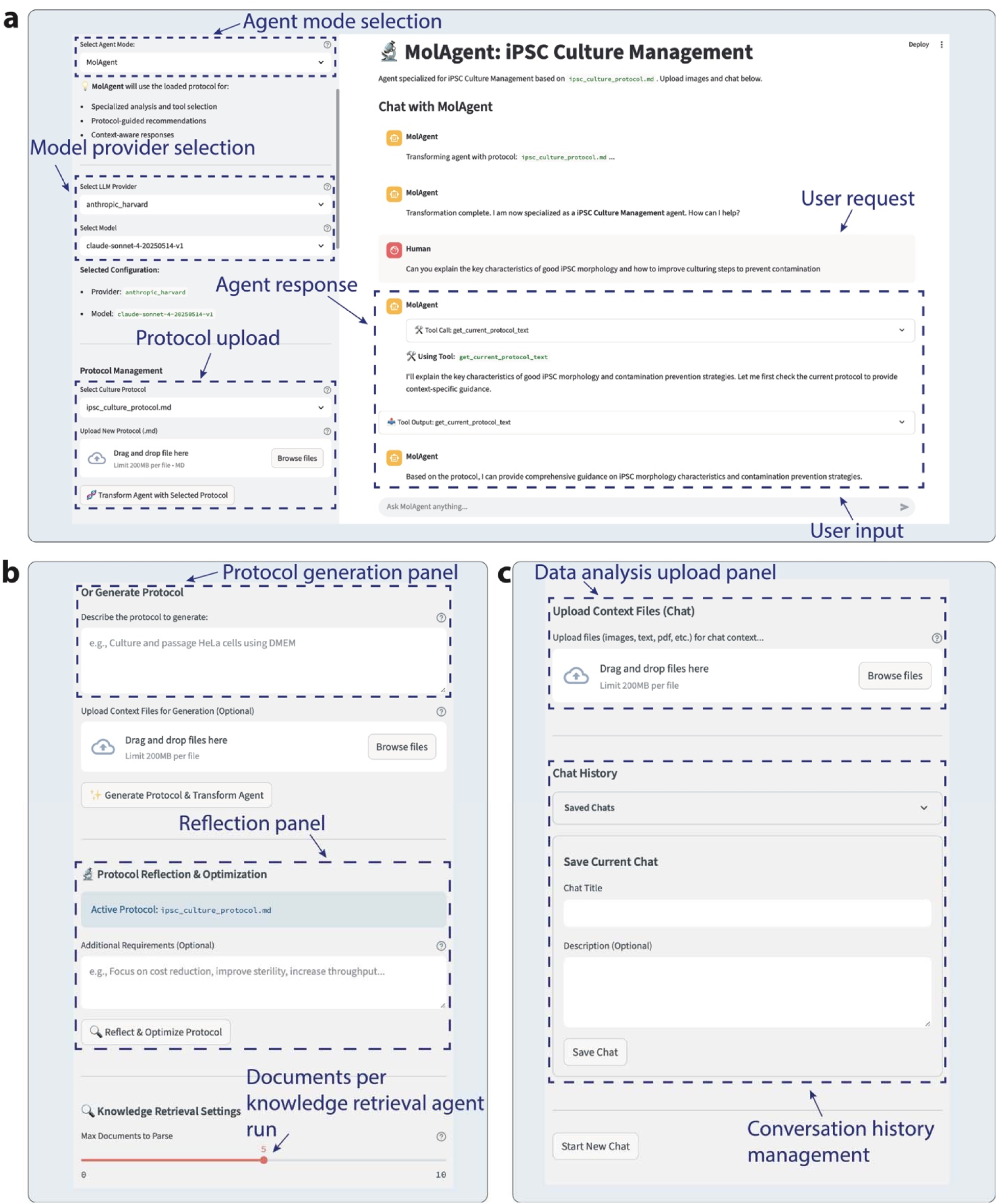
Frontend of MolAgent for cell and organoid research. Agentic Lab provides a web-based interface designed for conversational interactions in experimental design and analysis. Interface components include: **(a)**, agent mode and LLM core selection, together with the interaction panel where users submit natural language queries and receive context-aware responses; **(b)**, protocol generation and reflection panels that enable protocol creation, optimization, and refinement; and **(c)**, data upload and conversation history management panels for multimodal context input, analysis, and retrieval of past sessions. This interface offers researchers an accessible platform to design, optimize, and analyze experimental workflows without requiring advanced computational expertise.

**Extended Data Figure 3.**
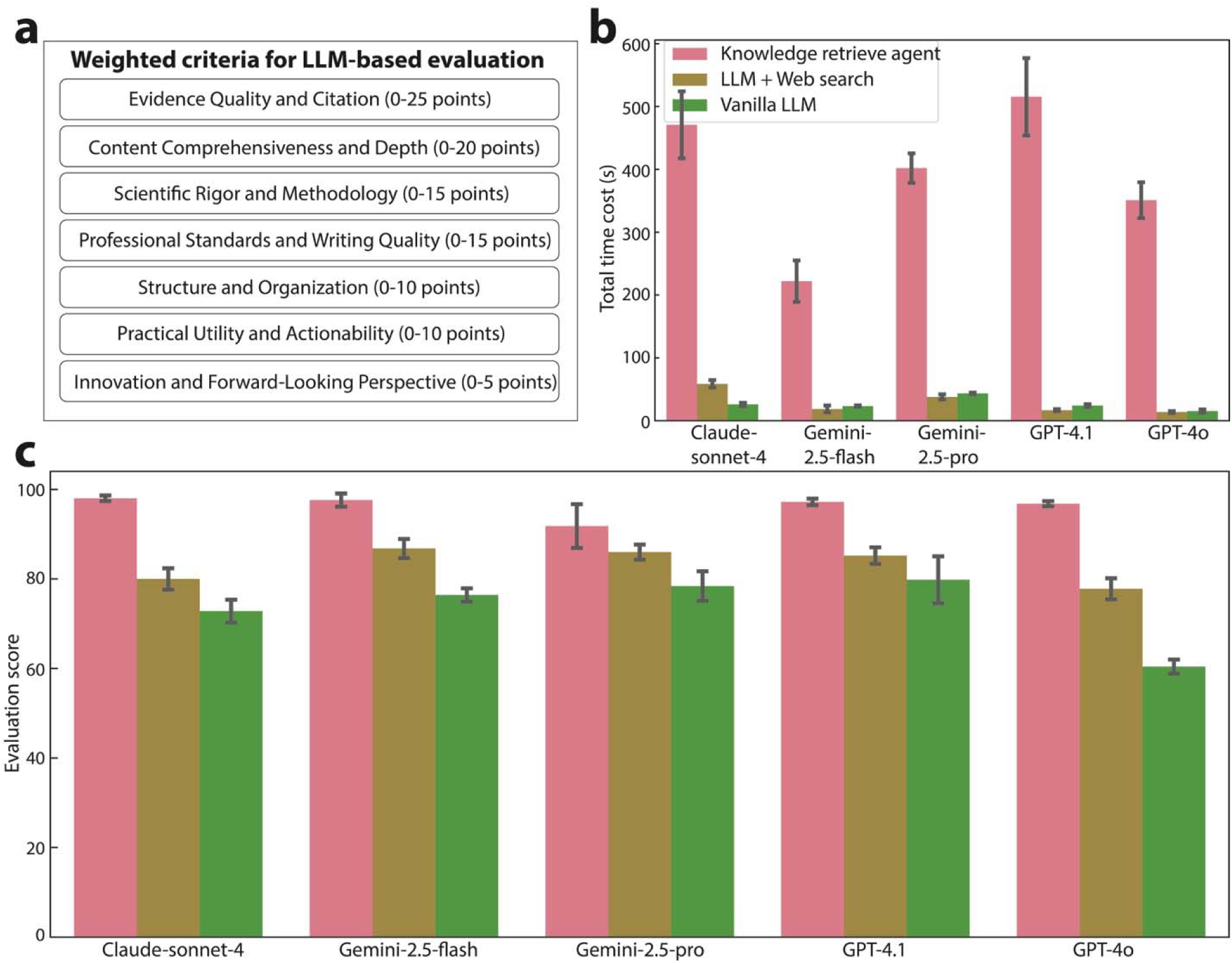
Benchmarking performance of the knowledge retrieval subagent. We compared the performance of vanilla LLMs (green), LLMs with web search capability (brown), and the knowledge retrieval subagent (pink) using a rule-based evaluation scheme. **(a)**, Weighted criteria used for evaluation, as determined by LLM-based scoring across different response dimensions. **(b)**, Time cost for generating responses across the three approaches. **(c)**, Overall evaluation scores across different LLMs, showing that the knowledge retrieval subagent consistently outperforms vanilla LLMs and LLMs with web search. n = 5 protocols of generation. Error bars represent mean ± SEM. across evaluation instances.

**Extended Figure 4.**
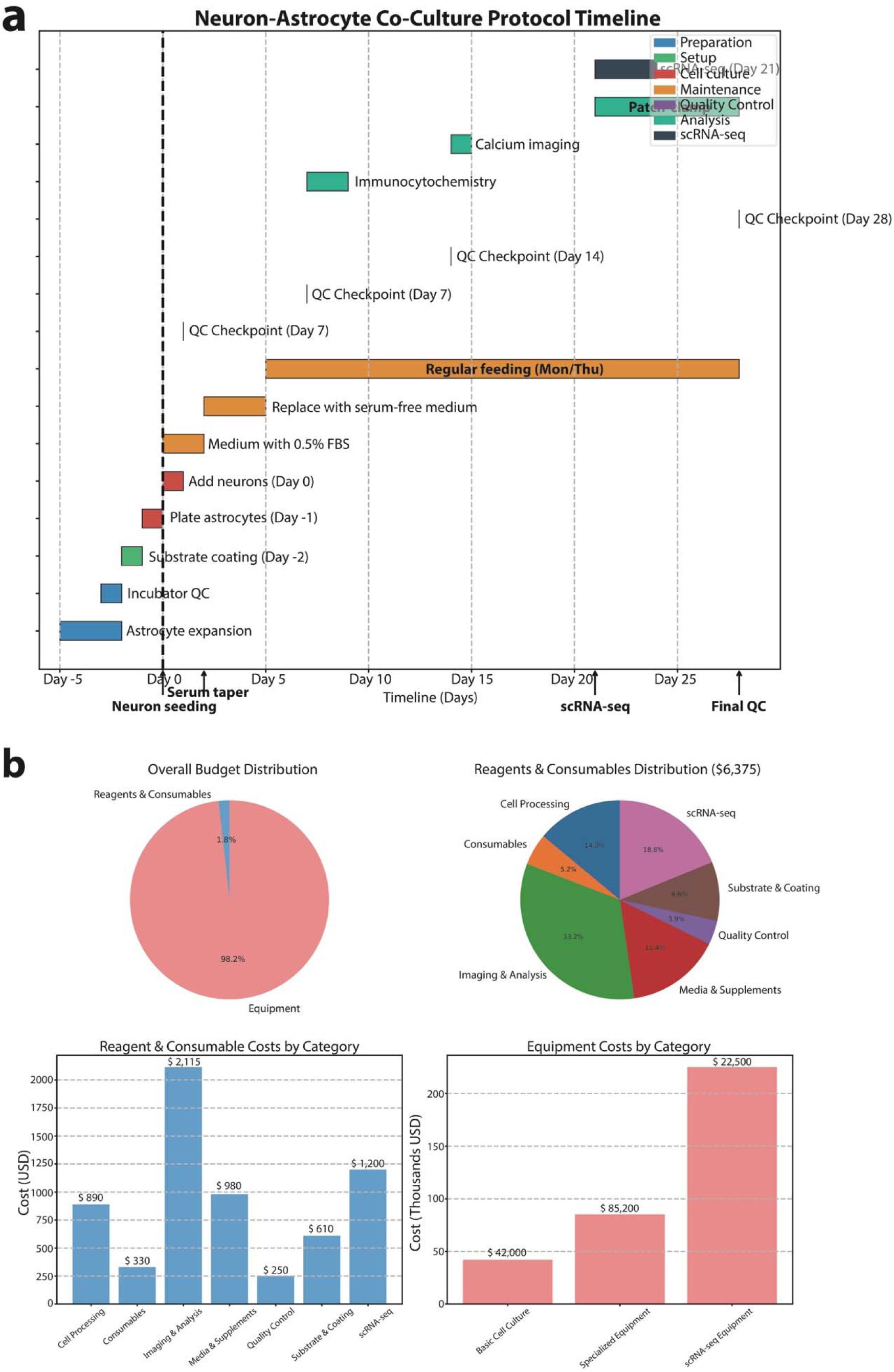
Example of plots for timeline and budget planning by Agentic Lab, relate to. figure 2**. (a)**, Gantt chart visualization of the neuron-astrocyte co-culture protocol generated. **(b)**, Visualization of experiment cost for the protocol in **(a)**.

**Extended Data Figure 5.**
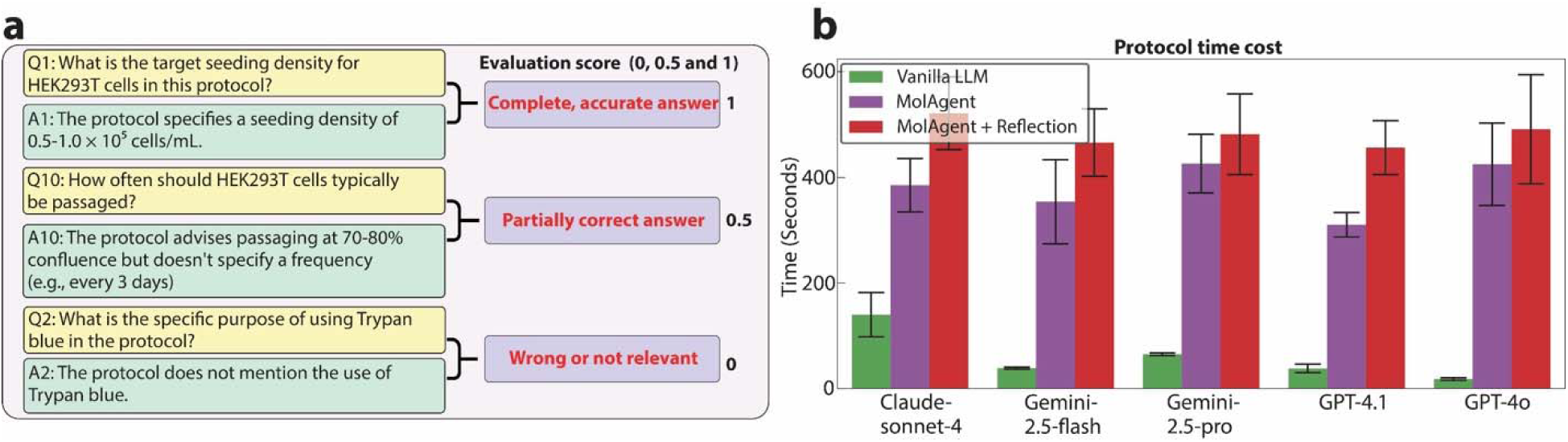
Benchmarking of protocol generation. (**a**), Grading scheme for Q&A evaluation, with examples of complete/accurate (score = 1), partially correct (score = 0.5), and incorrect/irrelevant (score = 0) answers. (**b**), Time cost of protocol generation for vanilla LLM (green), MolAgent (purple), and MolAgent with reflection (red). n = 47 Q&A pairs for each model. Error bars represent mean ± SEM.

**Extended Data Figure 6.**
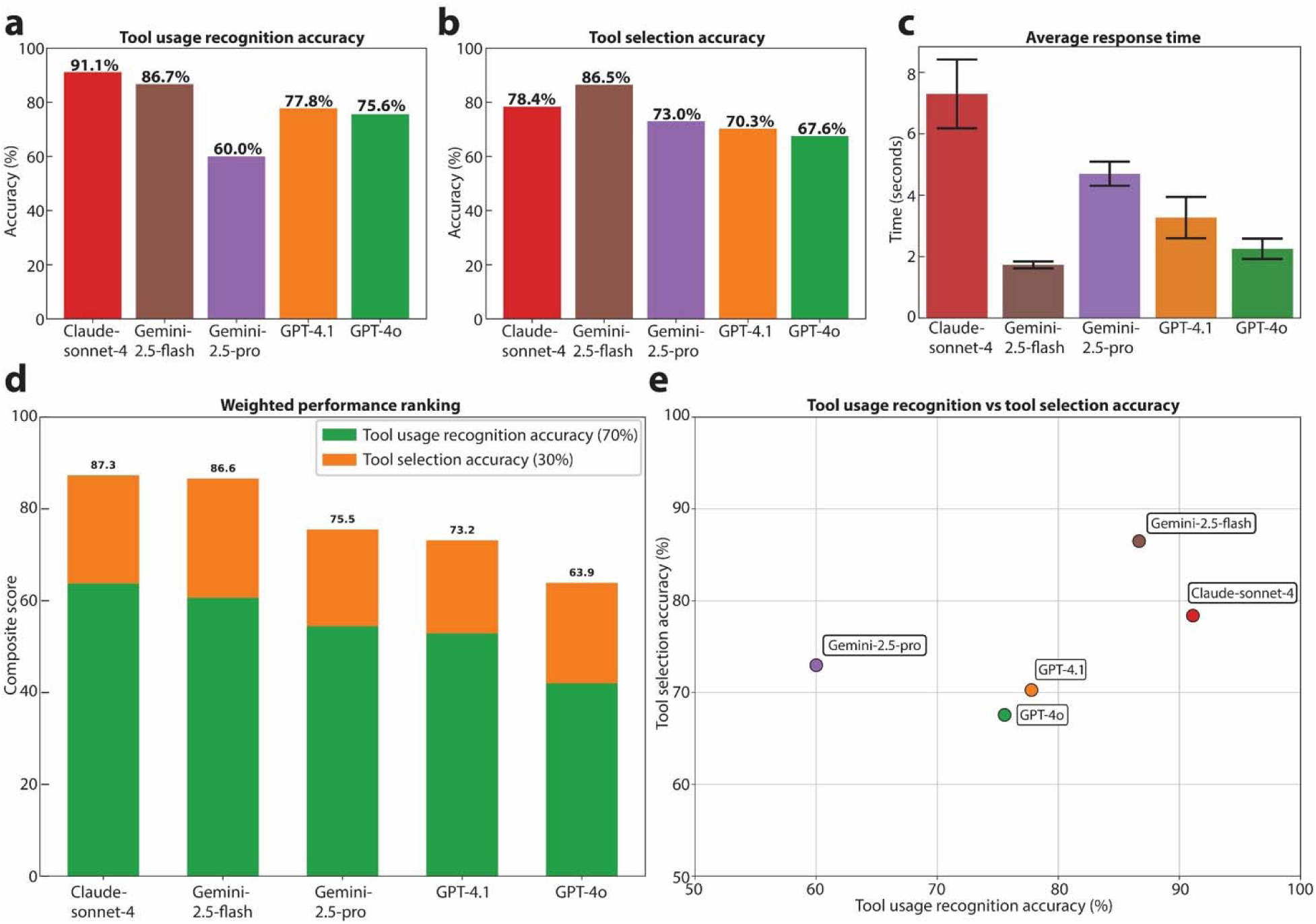
**Benchmarking tool adherence across LLM cores. (a-c)**, Two dimensions of tool adherence: tool usage recognition accuracy (determining whether a tool is needed) and tool selection accuracy (choosing the correct tool when required) are evaluated. **(a)**, Tool usage recognition accuracy for different LLMs. **(b)**, Tool selection accuracy. **(c)**, Average response time across models. Error bars represent mean ± SEM. n = 45 for each model. **(d)**, Weighted performance ranking combining 70% tool usage recognition accuracy and 30% tool selection accuracy. **(e)**, Two-dimensional scatter plot showing overall performance of different LLMs as agent cores. Results demonstrate variation in tool adherence across models.

**Extended Data Figure 7.**
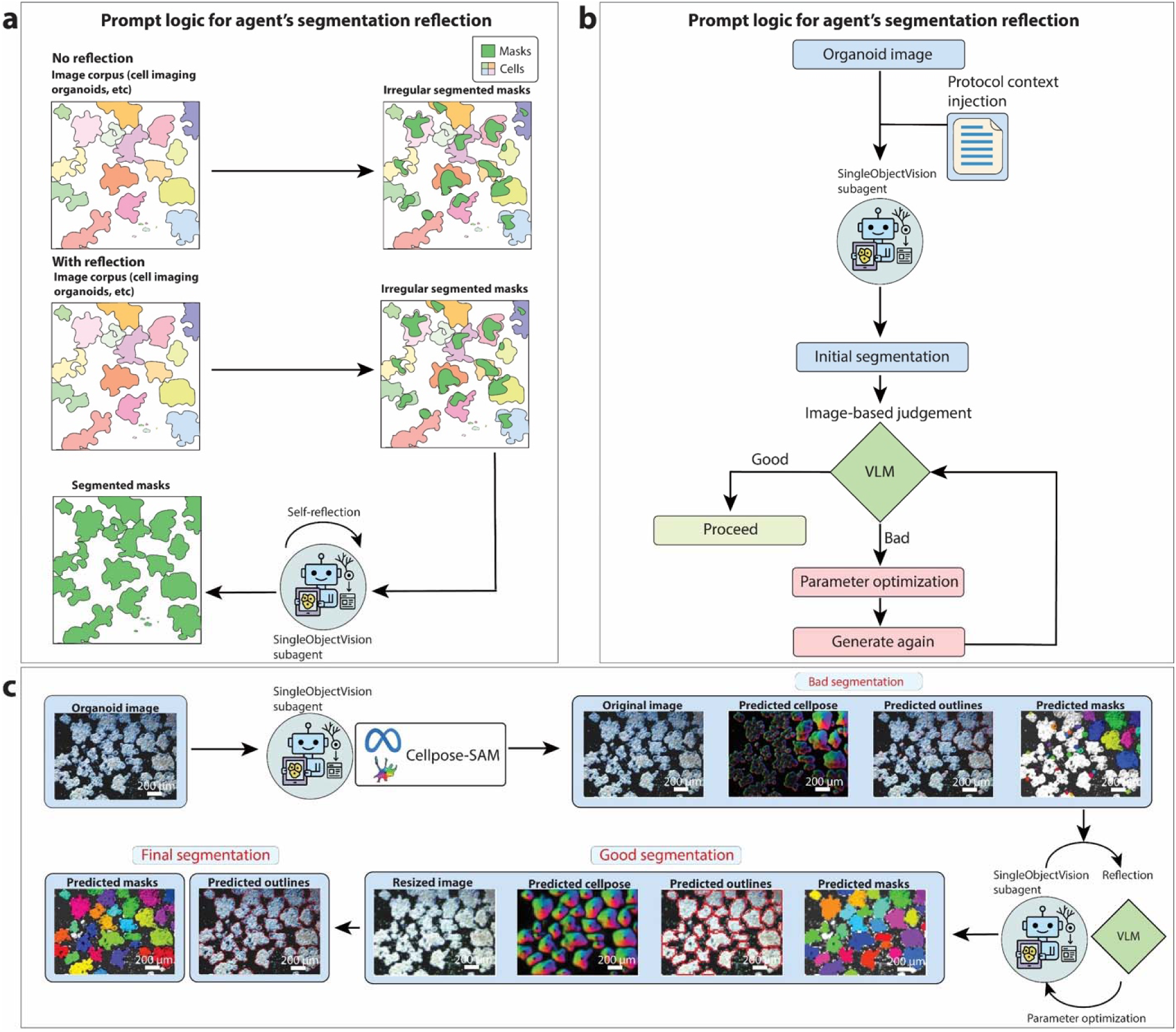
Reflection process of the SingleObjectVision subagent for segmentation optimization. **(a)**, Schematic of the agentic segmentation reflection pipeline. The agent leverages VLM and reasoning capabilities to evaluate the quality of generated masks, determine segmentation accuracy, and perform iterative parameter optimization until satisfactory masks are achieved. **(b)**, Prompt logic for the SingleObjectVision subagent showing how segmentation outputs are inspected, graded, and refined through reflection. **(c)**, Real-world example of the reflection process applied to an iPSC-derived pancreatic islet differentiation image at primitive gut tube stage. The agent integrates Cellpose-SAM segmentation with self-reflection to correct suboptimal outputs, leading to improved cell- and organoid-level masks. Scale bars, 200 μm.

**Extended Data Figure 8.**
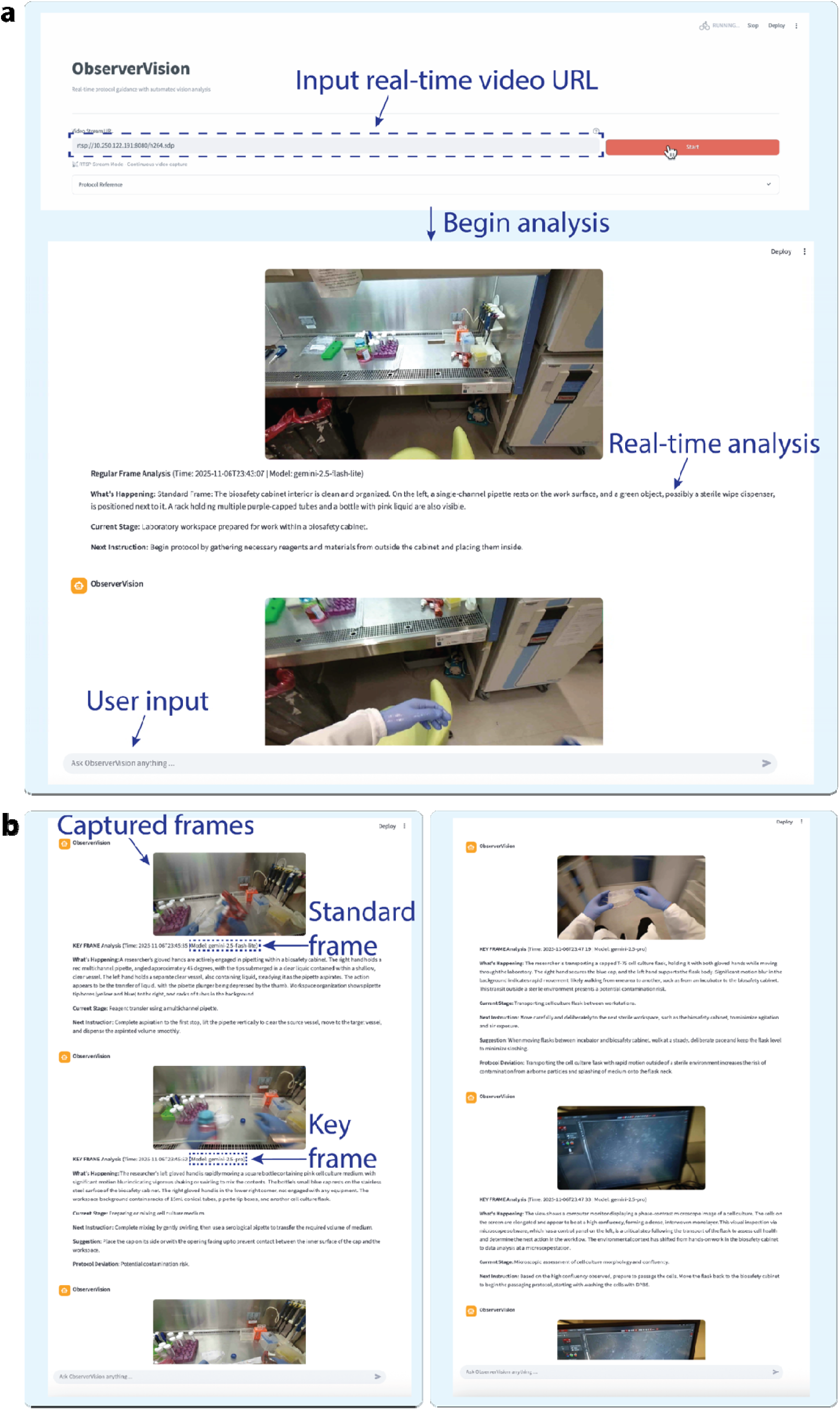
Frontend of physical AI system. **(a)** View from the human experimenter’s perspective at the laboratory bench, showing the Physical AI device in use. (**b**) User interface displayed on the AR glasses, featuring live video streaming, audio-to-text transcription, and interactive panels for protocol display and system-generated instructions and feedback.

**Extended Data Figure 9.**
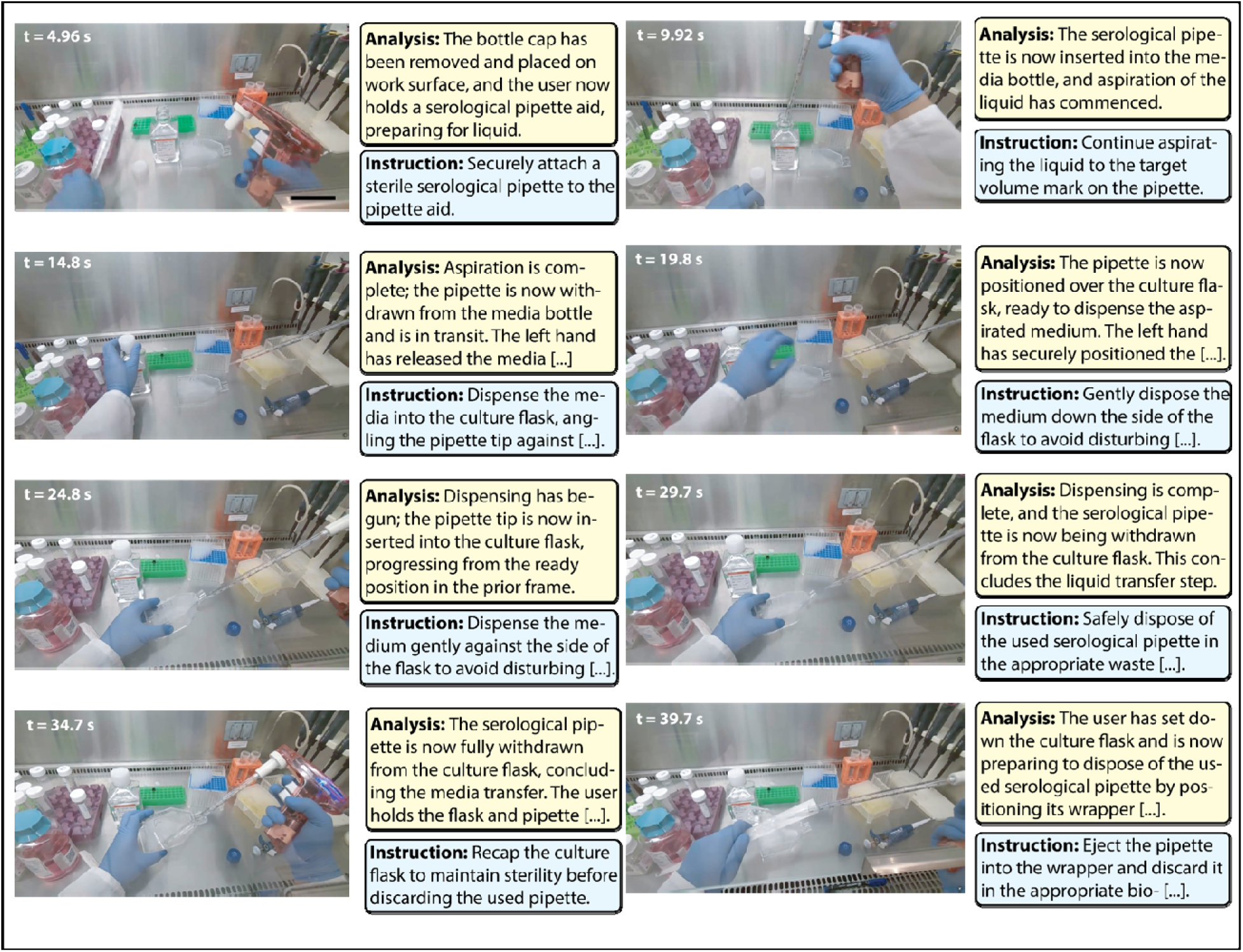
ObserverVision subagent-generated frame analysis and instruction at key frames for physical AI devices. Example key frames show automated recognition of user actions and generation of task-specific instructions, illustrating real-time perception, reasoning, and instruction generation enabled by the Agentic Lab system. Scale bar, 100 mm.

